# Multiple types of genomic variation contribute to adaptive traits in the mustelid subfamily Guloninae

**DOI:** 10.1101/2021.09.27.461651

**Authors:** Lorena Derežanin, Asta Blažytė, Pavel Dobrynin, David A. Duchêne, José Horacio Grau, Sungwon Jeon, Sergei Kliver, Klaus-Peter Koepfli, Dorina Meneghini, Michaela Preick, Andrey Tomarovsky, Azamat Totikov, Jörns Fickel, Daniel W. Förster

## Abstract

Species of the mustelid subfamily Guloninae inhabit diverse habitats on multiple continents, and occupy a variety of ecological niches. They differ in feeding ecologies, reproductive strategies and morphological adaptations. To identify candidate loci associated with adaptations to their respective environments, we generated a *de novo* assembly of the tayra (*Eira barbara)*, the earliest diverging species in the subfamily, and compared this with the genomes available for the wolverine (*Gulo gulo*) and the sable (*Martes zibellina*). Our comparative genomic analyses included searching for signs of positive selection, examining changes in gene family sizes, as well as searching for species-specific structural variants (SVs). Among candidate loci associated with phenotypic traits, we observed many related to diet, body condition and reproduction. For example, for the tayra, which has an atypical gulonine reproductive strategy of aseasonal breeding, we observe species-specific changes in many pregnancy-related genes. For the wolverine, a circumpolar hypercarnivore that must cope with seasonal food scarcity, we observed many changes in genes associated with diet and body condition. All types of genomic variation examined contributed substantially to the identification of candidate loci. This strongly argues for consideration of variation other than single nucleotide polymorphisms in comparative genomics studies aiming to identify loci of adaptive significance.

## Introduction

The Mustelidae are the most ecologically and taxonomically diverse family within the mammalian order Carnivora, representing a remarkable example of adaptive radiation among mammals that is rich with recent speciation events (Koepfli et al., 2008; Liu et al., 2020). Closely related mustelid species often inhabit vastly different ecosystems, where they experience diverse environmental challenges and are thus exposed to different evolutionary pressures. This is particularly pronounced in the mustelid subfamily Guloninae, within which species occupy a variety of ecological niches, ranging from scansorial omnivores in the neotropics to terrestrial hypercarnivores in circumpolar regions. Members of the Guloninae display a range of behavioural and physiological adaptations associated with environment-specific resource availability, and consequently differ markedly in feeding ecology, reproductive strategy and morphology (Heldstab et al., 2018; Zhou et al., 2011). Here, we focus on tayra, wolverine and sable (Figure 1), for which genomic resources are now available.

**Figure 1.**
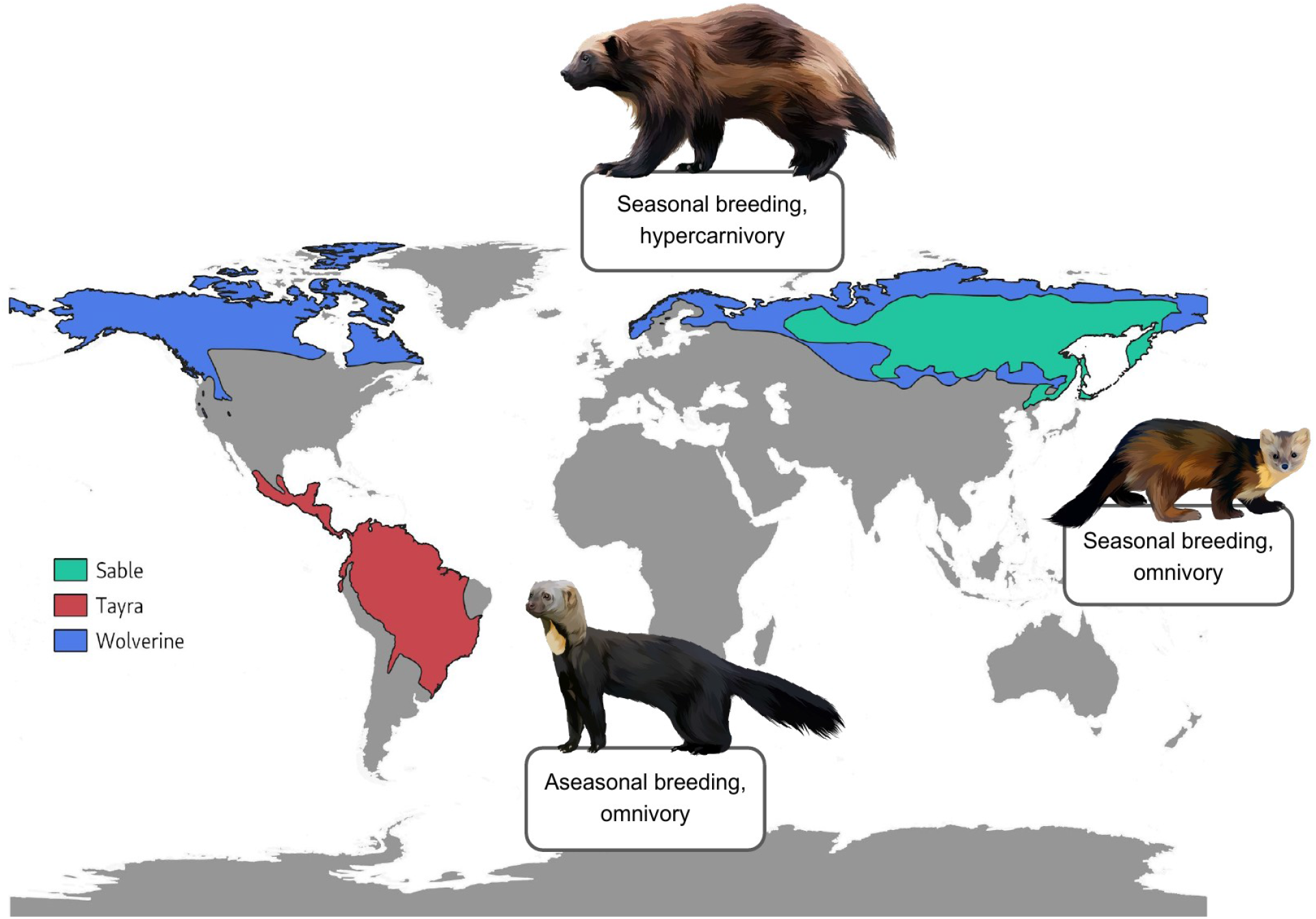
Distribution and species-specific traits of the tayra (*Eira barbara*), wolverine (*Gulo gulo*), and sable (*Martes zibellina*). Vector graphics of species are created based on royalty-free images (source: Shutterstock).

The tayra (*Eira barbara)* is a predominantly diurnal, solitary species that inhabits tropical and subtropical forests of Central and South America, ranging from Mexico to northern Argentina (Wilson & Mittermeier, 2009). It is a scansorial, opportunistic omnivore, feeding on fruits, small mammals, birds, reptiles, invertebrates and carrion. Caching of unripe fruit for later consumption has been observed (Soley & Alvarado-Díaz, 2011). Unlike other gulonine species, which are characterized by seasonal breeding and embryonic diapause, the tayra is an aseasonal polyestrous breeder and does not exhibit delayed implantation (Proulx & Aubry, 2017), which may be due to the less prominent seasonality and fluctuation in food availability in neotropical habitats (Heldstab et al., 2018).

The largest terrestrial mustelid, the wolverine (*Gulo gulo)*, is a circumpolar species, inhabiting alpine and boreal zones across North America and Eurasia (Ekblom et al., 2018). The wolverine is an opportunistic predator and facultative scavenger, either feeding on carrion or actively hunting medium to large-sized mammals, such as roe deer, wild sheep and occasionally moose (Pasitschniak-Arts & Larivière, 1995). Morphological and behavioural adaptations such as dense fur, plantigrade locomotion facilitating movement through deep snow, and food caching, enable wolverines to survive in cold habitats with limited food resources (Copeland & Kucera, 1997). In addition, wolverines occupy large home ranges, display territoriality, seasonal breeding and delayed implantation, traits indicating an adaptive response necessary for survival in scarce resource environments (Inman et al., 2012).

The sable (*Martes zibellina)* is distributed in the taiga and deciduous forests of north central and north eastern Eurasia. The sable is solitary and omnivorous, relying on hearing and olfaction to locate prey, even under a snow cover during winter months (Liu et al., 2020; Monakhov, 2011). Unlike wolverines, seasonal changes do not cause dramatic fluctuations in resource availability for sables as they are able to exploit a wider variety of food sources, and are adapted to tolerate short-term food scarcity (Mustonen et al., 2006). Their diet consists of small mammals, birds, nuts and berries, and in some instances food caching during the winter period has been reported (Monakhov, 2011). Similar to wolverines and many other species of mustelids, sables have a well-defined reproductive season and exhibit delayed blastocyst implantation (Proulx & Aubry, 2017).

To date, only a few studies have investigated adaptive variation in mustelids using comparative genomics (Abduriyim et al., 2019; Beichman et al., 2019; Liu et al., 2020; Miranda et al., 2021). Here, we generated a highly contiguous genome assembly of the tayra, an early diverging gulonine (Koepfli et al., 2008; Law et al., 2018), and compared it to previously published genomes of the wolverine and sable to identify the genetic basis underlying the adaptations to the diverse environments inhabited by these species.

In addition to identifying genes under positive selection, we investigated gene family evolution and structural variants (SVs), as these types of variants represent a significant source of intra- and interspecific genomic differentiation, affecting more nucleotides than single-nucleotide polymorphisms (Catanach et al., 2019). Gene copy number variation and large SVs can be associated with an adaptive response to new ecological circumstances (Rinker et al., 2019), and are thus an important source of genomic novelty to consider when studying adaptive divergence among species (Hecker et al., 2019). We focused on candidate loci linked to species-specific traits associated with response to environmental challenges, such as resource availability in the respective habitats of our study species.

## Materials and Methods

### Sequencing, Genome Assembly and Alignment

Whole blood from a captive male tayra was collected by the veterinary staff of the “Wildkatzenzentrum Felidae”, Barnim (Germany), during a routine medical checkup. High molecular weight (HMW) genomic DNA extraction was performed using the Qiagen MagAttract HMW DNA Kit, following the manufacturer’s protocol. We used 1ng of DNA and the Chromium Genome Reagents Kits Version 2 and the 10X Genomics Chromium Controller instrument with a microfluidic chip for library preparation. Sequencing was carried out on an Illumina NovaSeq 6000 (San Diego, CA, USA) with 300 cycles on an S1 lane.

We generated a *de novo* genome assembly using the 10x Genomics Supernova assembler v2.1.1 (Weisenfeld et al., 2017) with default parameters (assembly metrics given in SI Table S1). The assemblies of tayra, wolverine, sable and domestic ferret were assessed for gene completeness with BUSCO v4.1.2 using the mammalian lineage dataset mammalia_odb10 (Simão et al., 2015). To accurately identify repeat families, we used RepeatModeler v2 (Flynn et al., 2020) with the option “-LTRstruct” option, followed by Repeatmasker v4.1.2 (Smit, 2004) to identify and mask the modeled repeats in the tayra genome assembly.

For the purpose of demographic history and structural variation analyses, reads from all three Guloninae species were pre-processed (details given in Supplemental Information), and aligned to the domestic ferret (*Mustela putorius furo*) genome (MusPutFur1.0_HiC; Dudchenko et al., 2017, 2018; Peng et al., 2014) in local mode with Bowtie2 v2.3.5.1 (Langmead & Salzberg, 2012). Duplicated reads were removed with Picard Toolkit v2.23 (MarkDuplicates, Broad Institute 2019). Trimmed tayra reads were downsampled to ~38x with seqtk v1.3 (https://github.com/lh3/seqtk) prior to mapping to maintain uniformity among libraries and to avoid bias in variant calling.

Functional and biological roles of positively selected genes, loci affected by changes in gene family size, and structural variants, were explored using literature sources and online databases, including OrthoDB v10 (Kriventseva et al., 2019), Uniprot (The UniProt Consortium, 2017), and NCBI Entrez Gene (Maglott et al., 2011).

### Demographic reconstruction

Analysis of demographic history for tayra and sable was performed with PSMC v0.6.5 (Li & Durbin, 2011) using the following parameters (repeated 100 times for bootstrapping): *psmc -N25 -t15 -r5 -b -p ‘4+25*2+4+6’ -o round-${ARRAY_TASK_ID}.psmc ${name}.split.psmcfa*. Results for each genome were plotted with psmc_plot.pl, and the mutation rate was set to 1e-08 substitutions per site per generation (Cahill et al., 2016; Dobrynin et al., 2015). Generation times were set to 7.4 years for tayra, 5.7 for sable (Pacifici et al., 2013) and 6 years for wolverine (Ekblom et al., 2018).

### Reference-based scaffolding

Using the domestic ferret genome as reference, we generated pseudochromosome assemblies for tayra, wolverine and sable, to visualize heterozygosity along chromosomes. Scaffolding was performed using RaGOO v1.1 (Alonge et al., 2019). The X chromosome in the domestic ferret assembly was identified via whole genome alignment to the domestic cat (*Felis catus*) Felis_catus_9.0 assembly (Buckley et al., 2020) and ZooFISH data available from the *Atlas of Mammalian Chromosomes* (Cavagna et al., 2000; O’Brien et al., 2020). Whole genome alignment was performed using LAST v971 (Frith & Kawaguchi, 2015).

Variant calling followed by quality filtration was performed using the BCFtools pipeline v1.10 (Poplin et al., 2018). Low-quality variants were removed (BCFtools filter, ‘QUAL < 20.0 || (FORMAT/SP > 60.0 || FORMAT/DP < 5.0 || FORMAT/GQ < 20.0)’). In each sample, positions with coverage lower or higher than 50-250% of the whole genome median value were removed. Of the remaining positions only those common to all samples were retained. Finally, SNPs with uncalled genotypes in any sample and variants with the same genotypes for all samples were removed. For visualization, heterozygous SNPs were counted in non-overlapping sliding windows of 1 Mbp (counts scaled to SNPs per kbp). Indels were not included due to the low quality of calls from short reads. SNP density plots were created using the MACE package (https://github.com/mahajrod/MACE).

### Phylogenomic data preparation, analysis and dating

We performed sequence alignments and filtering of highly divergent alignment segments for 6020 coding genomic regions of single-copy orthologs shared across eight species of carnivores, using the software MACSE v2 (Ranwez et al., 2011). Alignments were further trimmed for highly divergent regions, excessive missing data, and substitution model adequacy (further details in SI). Gene trees were estimated from gene regions by first selecting the best substitution model from the GTR+F+Γ+I+R family (Kalyaanamoorthy et al., 2017), and calculating approximate likelihood-ratio test (aLRT) branch supports (Anisimova & Gascuel, 2006), as implemented in IQ-TREE v2 (Minh et al., 2020b).

Species tree estimates were performed using (i) concatenated sequence alignments for maximum likelihood inference using IQ-TREE v2, and (ii) gene trees for inference under the multi-species coalescent using the summary coalescent method in ASTRAL-III (Zhang et al., 2018). The maximum likelihood estimate of the species tree was accompanied by aLRT branch supports, while summary coalescent inference was accompanied by local posterior probabilities (Sayyari & Mirarab, 2016). The decisiveness of the data regarding the phylogenetic signals was examined using gene- and site-concordance factors, calculated in IQ-TREE v2 (Minh et al., 2020a).

Bayesian molecular dating analysis was performed using MCMCtree in PAML v4.8 (Yang, 2007), assuming the reconstructed species tree from ASTRAL-III and using the genomic regions with gene trees concordant with the species tree (see SI for further details). Regions were partitioned by codon positions, each modelled under individual GTR+Γ substitution models. We used an uncorrelated gamma prior on rates across lineages and a birth-death prior for divergence times. Fossil calibrations are listed in the SI. The posterior distribution was sampled every 1×10^3^ MCMC steps over 1×10^7^ steps, after a burn-in phase of 1×10^6^ steps. We verified convergence to the stationary distribution by comparing the results from two independent runs, and confirming that the effective sample sizes for all parameters were above 1000 using the R package coda (Plummer et al., 2006).

### Positive selection on single-copy orthologs

To investigate genes under positive selection, the CDS corresponding to 1:1 orthologs were aligned for the eight aforementioned carnivoran species. Multiple sequence alignments (MSAs) were constructed with PRANK v120716 (Löytynoja, 2014), and 17 MSAs were removed due to short alignment length. The CODEML module in the PAML v4.5 package was used to estimate the ratio of non-synonymous to synonymous substitutions, also called *d_N_* /*d_S_* or *ω* (Yang, 2007). We applied the one-ratio model to estimate the general selective pressure acting among all species, allowing only a single *d_N_*/*d_S_* ratio for all branches. A free-ratio model was also used to estimate the *d_N_*/*d_S_* ratio of each branch. Furthermore, the CODEML branch-site test for positive selection was performed on 6003 ortholog alignments for three separate foreground branches: *Eira barbara*, *Gulo gulo* and *Martes zibellina* (Zhang et al., 2005). Statistical significance was assessed using likelihood ratio tests (LRT) with a conservative 10% false discovery rate (FDR) criterion (Nielsen et al., 2005). Orthologs with a free-ratio > 2 in the branch model were considered for further analysis of signatures of positive selection.

To account for differences in genome assembly quality, we evaluated the alignments of selected orthologs based on the transitive consistency score (TCS), an extension to the T-Coffee scoring scheme used to determine the most accurate positions in MSAs (Chang et al., 2014). Additionally, alignments were visually inspected for potential low-scoring MSA portions.

### Gene family evolution

To investigate changes in gene family sizes, we constructed a matrix containing 7838 orthologs present as either complete ‘single-copy’, complete ‘duplicated’, or ‘missing’, identified using the BUSCO genome assembly completeness assessment of all eight carnivoran genomes. Orthologs were retained if they were detected in at least four species (including *Felis catus* as an outgroup) to obtain meaningful likelihood scores for the global birth and death (*λ*) parameter.

We applied a probabilistic global birth and death rate model of CAFE v4.2.1. (Han et al., 2013) to analyse gene gains (‘birth’) and losses (‘death’) accounting for phylogenetic history. First, we estimated the error distribution in our dataset, as genome assembly and annotation errors can result in biased estimates of the average rate of change (*λ*), potentially leading to an overestimation of *λ*. Following the error distribution modelling, we ran the CAFE analysis guided by the ultrametric tree estimated earlier, calculating a single *λ* parameter for the whole species tree. The CAFE results were summarized (SI Table 4A) with the python script *cafetutorial_report_analysis.py* (https://github.com/hahnlab/CAFE).

### Structural variation

We applied an ensemble approach for structural variant (SV) calling, encompassing three SV callers: Manta v1.6.0 (Chen et al., 2016), Whamg v1.7.0 (Kronenberg et al., 2015) and Lumpy v0.2.13 (Layer et al., 2014). SV calls originating from reads mapping in low complexity regions and withpoor mapping quality were removed from all three call sets. We retained Manta calls with paired-read (PR) and split-read (SR) support of PR>=3 and SR>=3, respectively. To reduce the number of false positive calls, the Whamg call set was filtered for potential translocation events, as Whamg flags but does not specifically call translocations. We further removed calls with a low number of reads supporting the variant (PR, SR) from the Whamg (A < 10) and the Lumpy call set (SU < 10). All SV call sets were filtered based on genotype quality (GQ ≥ 30). Whamg and Lumpy SV call sets were genotyped with Svtyper v0.7.1 (Chiang et al., 2015) prior to filtering. Only scaffolds assigned to chromosomes were included in further analyses. Survivor v1.0.7 (Jeffares et al., 2017) was used to merge and compare SV call sets within and among samples. The union set of SV calls among the three gulonine species containing species-specific and shared variants was annotated, using Liftoff v1.5.1 (Shumate & Salzberg, 2020) for preparation of reference genome annotation, and Ensembl Variant Effect Predictor v101.0 (McLaren et al., 2016) for identifying variants affecting protein-coding genes. Gene ontology analysis was performed with Shiny GO (Ge et al., 2020) with an FDR < 0.05 for each SV type (excluding inversions) overlapping multiple protein-coding genes (> 5 genes).

## Results

### Genome Assembly

We generated a highly contiguous reference genome assembly for the tayra (*Eira barbara*). Extracted genomic DNA had an average molecule size of 50.75 kb and was sequenced to ~76-fold coverage (SI Table S1). The final assembly showed a total length of ~2.44 Gb (excluding scaffolds shorter than 5 kb), with a contig N50 of 290 kb, scaffold N50 of 42.1 Mb, and identity in 95% of all positions in an alignment with domestic ferret genome (SI Figure S1). The tayra assembly has higher contiguity than the Illumina-only based assemblies of both wolverine and sable, but it is more fragmented than the chromosome-length domestic ferret (*Mustela putorius furo)* assembly (Table 1, SI Figure S2A) that we used as reference genome for some analyses. The haploid tayra genome of ~2.4 Gb is contained in 162 scaffolds (>100 kb) with 40 scaffolds having a length above 50 Mb (SI Figure S2A).

**Table 1.**
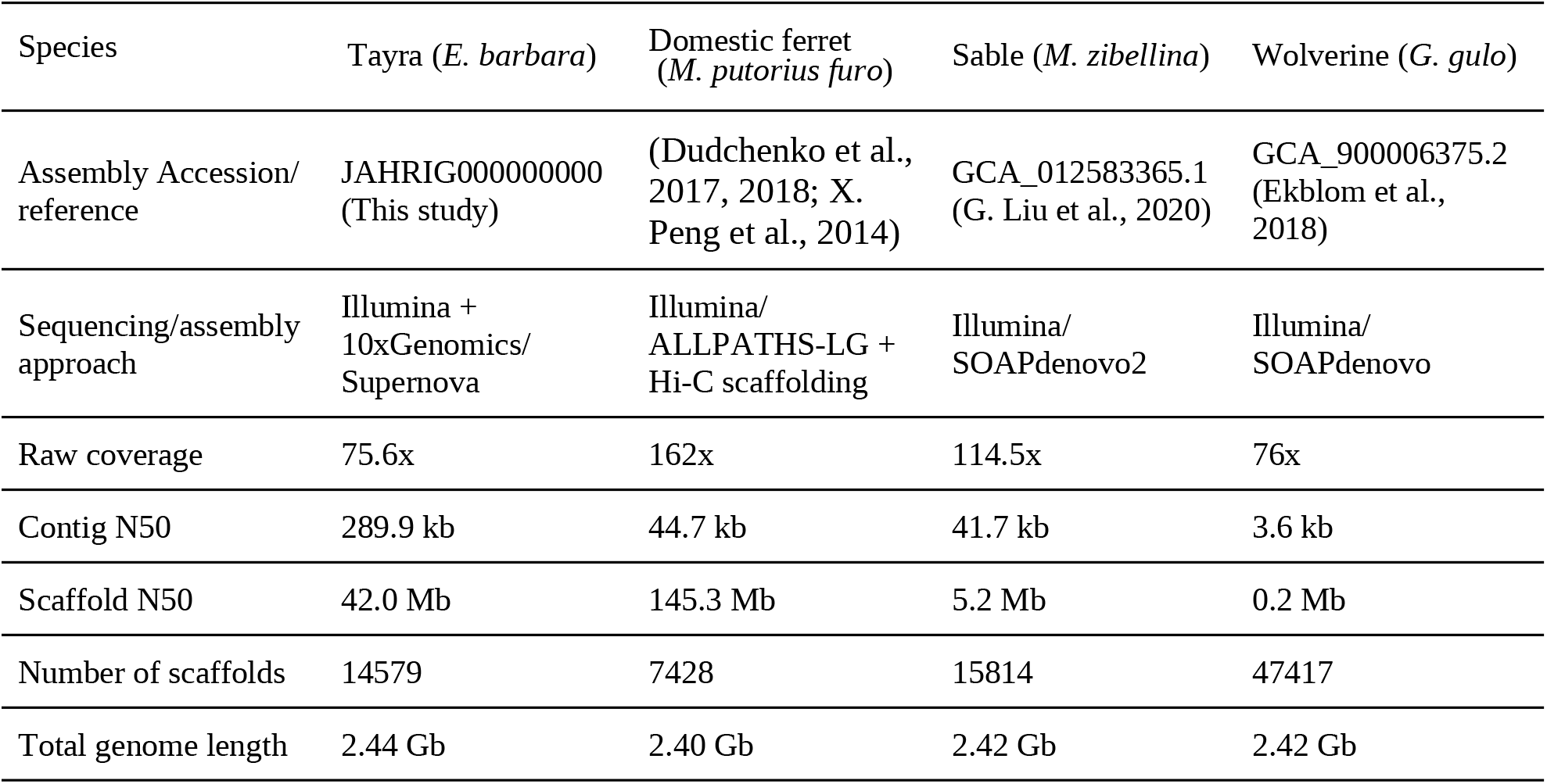
Comparison of genome assembly metrics among four mustelid species.

The tayra assembly has high gene completeness as assessed with BUSCO v4.1.2 using 9226 conserved mammalian orthologs. In total, 8540 (92.5%) complete Benchmarking Universal Single-Copy Orthologs (BUSCOs) were identified, encompassing 8492 (92.0%) of complete and single-copy, and 48 (0.5%) complete and duplicated orthologs. Additionally, 104 (1.1%) orthologs were fragmented and 582 (6.4%) were missing. As measured by this metric, the tayra genome has higher gene completeness than the published genomes of wolverine, sable or domestic ferret (SI Figure S2B).

### Repetitive elements

The repeat landscape of the tayra assembly contains ~0.85 Gb of repetitive elements (SI Table S2). L1 type LINE elements are the most abundant, constituting 23% of the tayra genome. L1 elements also show signs of recent proliferation in comparison to DNA transposons and LTR retroelements (SI Figure S3). Endogenous retroviruses constitute 3.8% of the tayra genome and can be classified as Gammaretroviruses and Betaretroviruses.

The overall repeat landscape of the tayra genome assembly is comparable to other carnivore genomes (Liu et al., 2020; Peng et al., 2018). It is similar to that of the sable genome, differing mostly in the number of L1 LINE elements, which have been recently proliferating and accumulating within the tayra genome more than in other Guloninae genomes. The diversity of endogenous retroviruses is similar to other mustelids. Although endogenous delta-retroviruses have been described from a broad range of mammal genomes, including several smaller carnivores like mongoose (family Herpestidae) and the fossa (*Cryptoprocta ferox*) (Hron et al., 2019), no delta-retroviruses were found in the genome of tayra.

### Demographic reconstruction

Reconstruction of historical demography for tayra and sable using the Pairwise Sequentially Markovian Coalescent model (PSMC; Li & Durbin, 2011) showed similar effective population sizes (Ne) during the last one million years of the Pleistocene, but then start to differ around twenty thousand years ago, when the Ne of the sable increases relative to that of the tayra (SI Figure S4). Historical demography for the wolverine was reconstructed from Ekblom et al. 2018, and shows the lowest Ne among the three gulonine species.

### Nucleotide diversity

The tayra, sable and wolverine assemblies were generated using different approaches and differ significantly in contiguity (Table 1). To address this issue, we generated pseudochromosome assemblies for each species using the chromosome-length assembly of the domestic ferret as reference. The domestic ferret has more chromosomes than the other species (2n=40 vs. 2n=38), and the same number (20) of pseudochromosomes (Lewin et al., 2019) were obtained after scaffolding in each case. For each assembly, we identified the X chromosome (labeled as *ps_chrX*) and arranged pseudoautosomes (labeled as *ps_aut1 - ps_aut19*) according to the length of the original scaffolds in the domestic ferret reference. This allowed us to verify the sex of the animals using a coverage-based approach (SI Figure S5), which confirmed morphological sexing for the tayra (male) and wolverine (female) individuals. While the sable individual is referred to as a male (Liu et al., 2020), our analysis suggests it is a female (further details given in SI).

We counted heterozygous SNPs in 1 Mbp stacking windows for all three species and scaled it to SNPs per kbp (Figure 2). Median values for tayra, sable and wolverine were 1.89, 1.44 and 0.28 SNPs per kbp, respectively, the latter being in agreement with previous findings (Ekblom et al., 2018). All scaffolds of ≥ 1 Mbp in pseudochromosome assemblies were taken into account. Exclusion of ps_chrX resulted in slight increases of medians to 1.93, 1.47 and 0.29, respectively (SI Table S3, SI Figure S6). Regardless of whether the ps_chrX was included or excluded, the genome-wide diversities among the three species were significantly different (*p*-values << 0.001, Mann-Whitney test).

**Figure 2.**
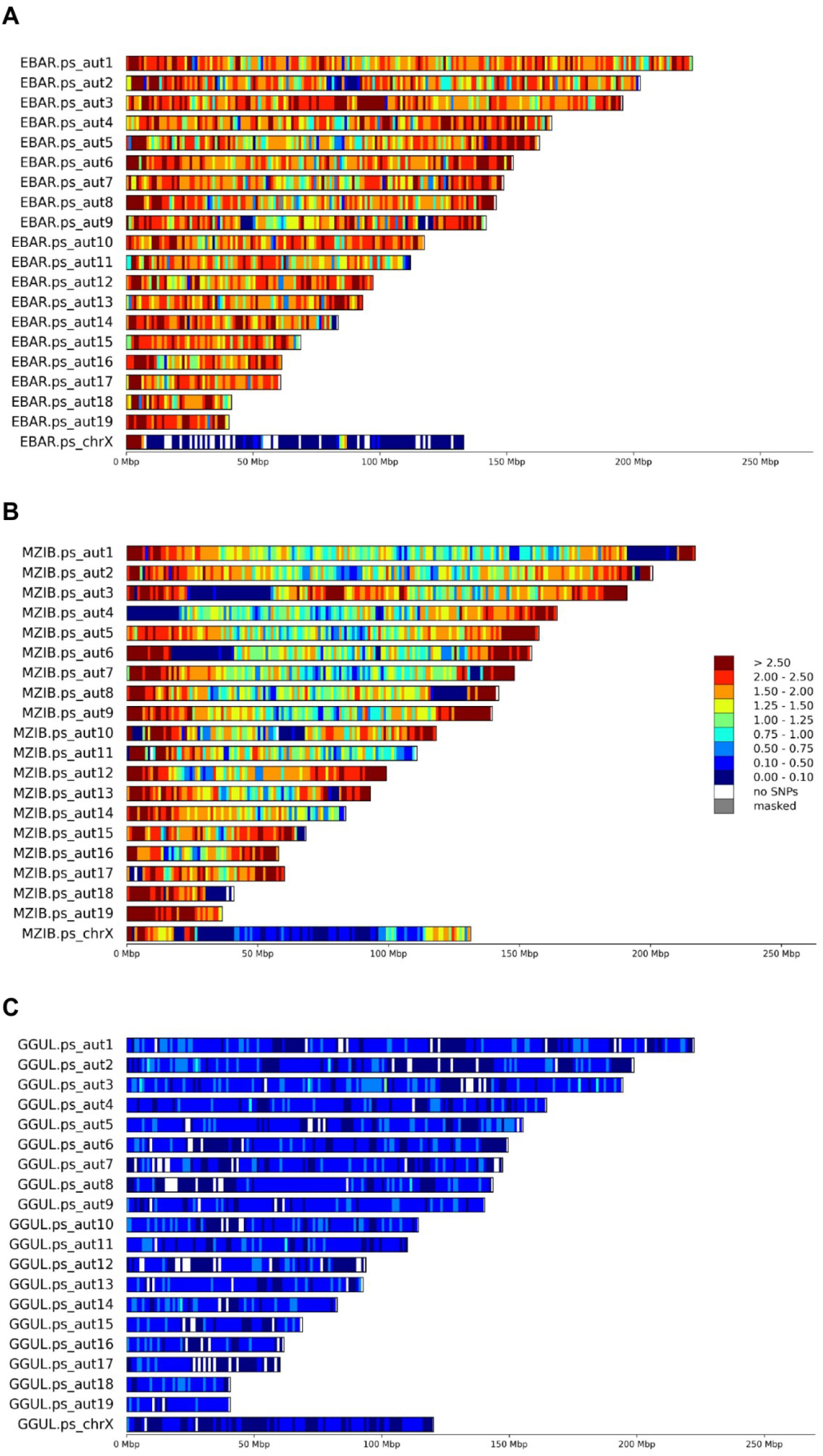
Heterozygosity density among pseudochromosomes for (A) tayra, (B) sable and (C) wolverine. Heterozygous SNPs were counted in stacking windows of 1 Mbp and scaled to SNPs per kbp. Tayra is a male individual and thus heterozygous SNP density is underestimated (due to only one X chromosome), while sable and wolverine are females and therefore likely representative of true SNP density.

### Phylogenomics and molecular dating

We reconstructed the phylogenetic relationships of the following carnivoran species: domestic cat, domestic dog, walrus, northern elephant seal, domestic ferret, wolverine, sable, and tayra. Phylogenomic analyses using concatenation and summary coalescent methods led to an identical resolution of the relationships among mustelid taxa. Within Guloninae, the wolverine and sable were placed as sisters, to the exclusion of the tayra (Figure 3). Branch supports were maximal across all branches using both approximate likelihood-ratio test (aLRT) and local posterior probabilities. Similarly, concordance factors (CF) for genes and sites (gCF, sCF) were high across branches and consistently more than twice as high as the values of the discordance factors (gDF, sDF). The lowest concordance factors were those in support of the resolution of *Gulo* and *Martes* as sisters (gCF = 64.52, sCF = 54.38). However, the discordance factors were less than half these values (gDF < 14, sCF < 24), suggesting substantial decisiveness across genes and sites for this resolution.

**Figure 3.**
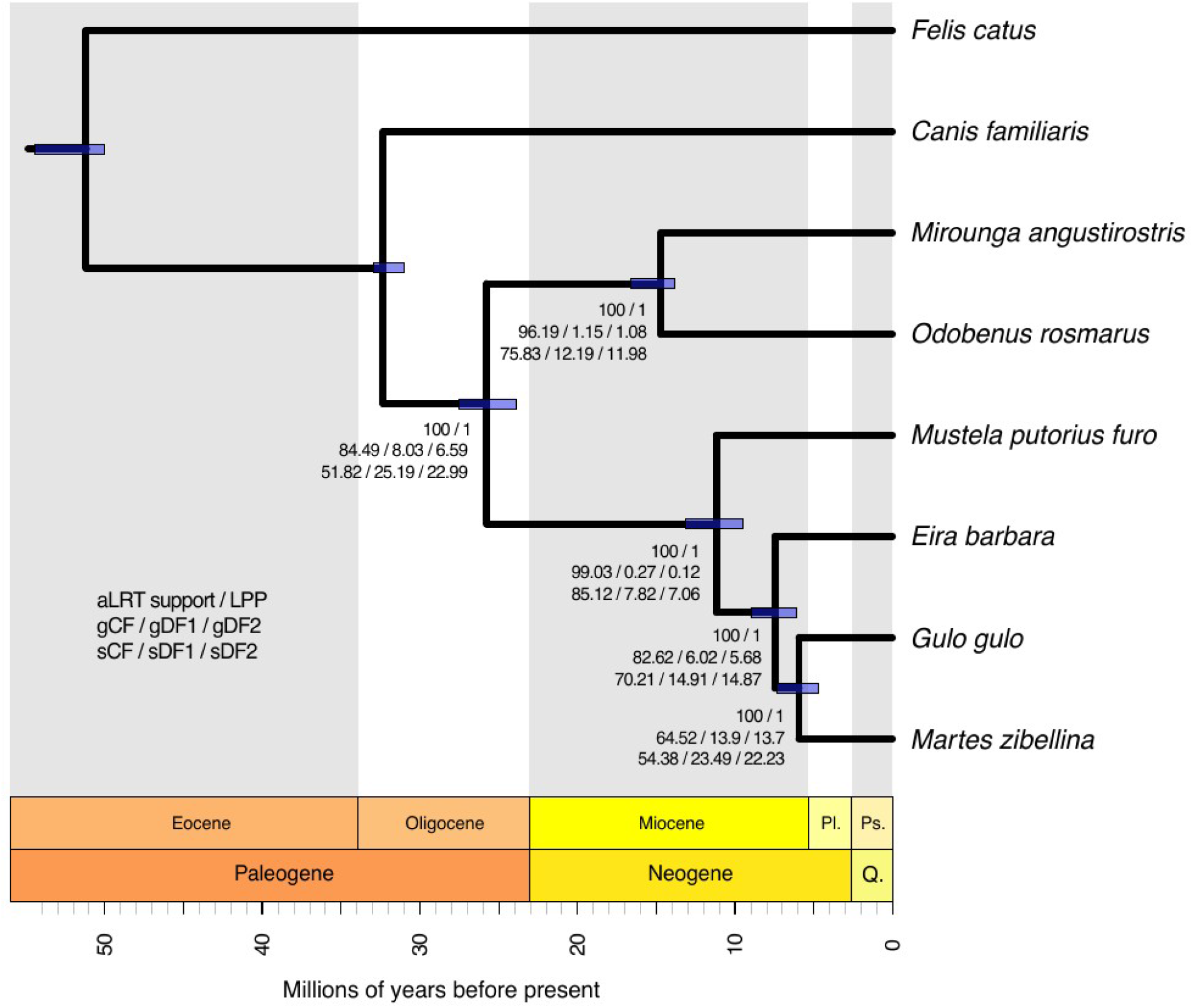
Phylogenetic tree and divergence times of Guloninae and five other carnivorans. The mean age of each node is shown, with 95% confidence intervals depicted as purple bars. The gene and site concordance (gCF, sCF) and discordance (gDF, sDF) factors are given, along with the branch support for both approximate likelihood-ratio test (aLRT) and local posterior probabilities (LPP).

Molecular dating was performed using 992 single-copy orthologous gene regions (comprising 0.53 million sites). Divergence time estimates across mustelids were largely in agreement with previous findings (Koepfli et al., 2008; Law et al., 2018; Li et al., 2014; Sato et al., 2012), placing the split between *Mustela* and Guloninae at 11.2 million years ago (Mya) (highest posterior density interval (HPDI) between 13.1 and 9.5 Mya), and the split between *Eira* and the *Gulo-Martes* group at 7.5 Mya (HPDI between 9 and 6.1 Mya). The split between *Gulo* and *Martes* was dated at 5.9 Mya (HPDI between 7.4 and 4.7 Mya).

### Positive selection on single-copy orthologs

In the three Guloninae species, we found sites under positive selection (5 > dN/dS ratio > 1, (Barnett et al., 2020) in 55 single-copy orthologs that were highly significant (free-ratio > 2). Of these 55 positively selected genes (PSGs), 15 were observed in tayra, 22 in wolverine, and 18 in sable (Figure 4A-B). Gene names, descriptions and functions are given in SI Table S4.

**Figure 4.**
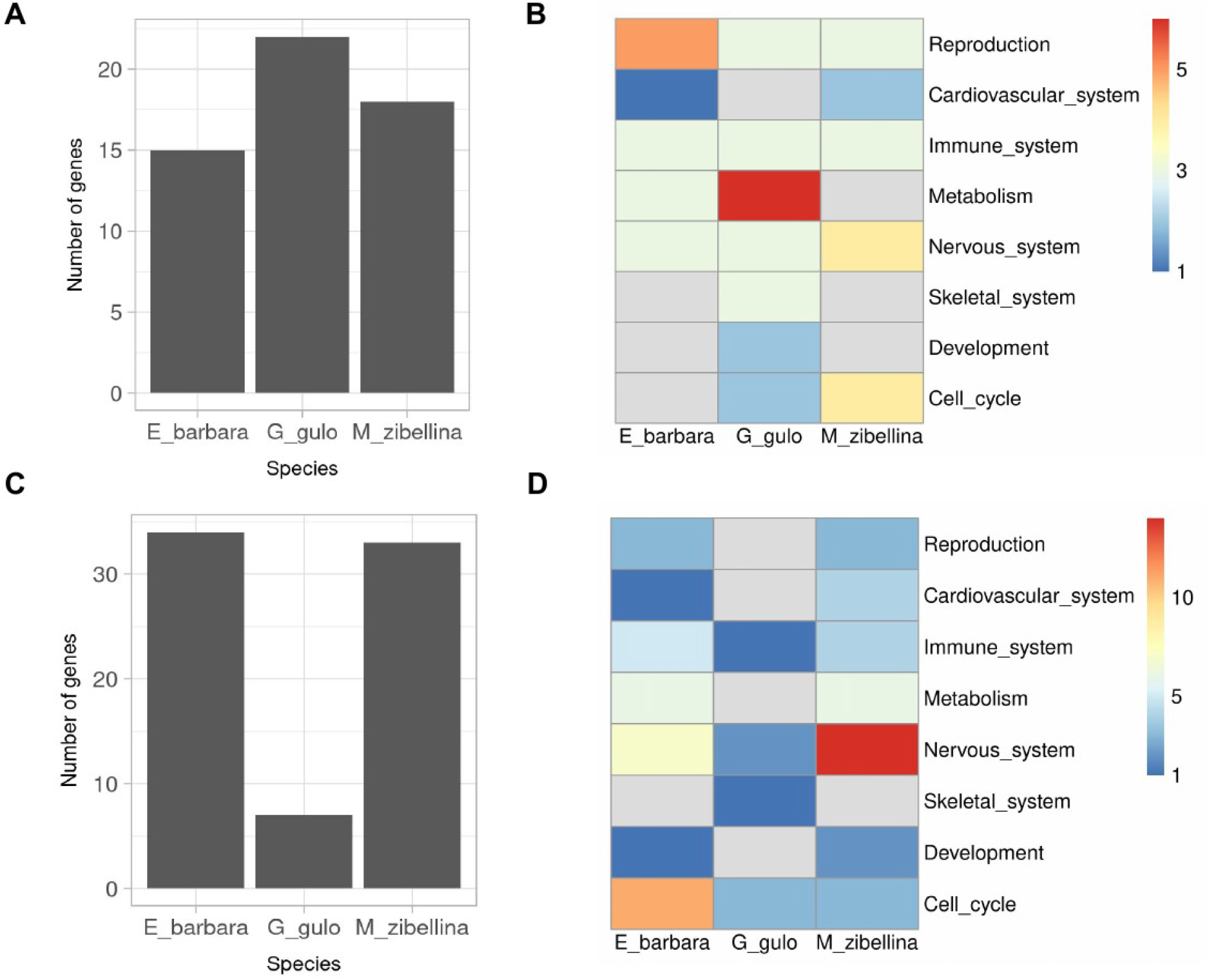
Number of candidate genes and their functional groups. Genes identified from analyses of (A,B) positive selection on single genes (PSG), and (C,D) gene family expansions. Heatmap scale represents the number of genes. Heatmap cells in grey color indicate no observations for a given variable.

Among the 15 PSGs we detected in tayra, five are associated with reproduction (*NSMCE1*, *ETV2*, *SPATA25*, *MUC15*, and *PIH1D2*) with functions involving spermatogenesis, placenta and embryo development, and blood vessel morphogenesis. Among the remaining ten PSGs, *HSPB6* is involved in vasodilation and muscle contraction, *DERA* is associated with environmental stressors, including exposure to toxins, and uricase (*UOX*) is a liver enzyme involved in purine catabolism and regulation of fructose metabolism. Three PSGs (*IP6K3*, *MAGIX* and *FAM149B1*) are found to be associated with the nervous system, synapse formation and structural plasticity, as well as motor skills and coordination. Three further PSGs (*DUSP19*, *TNLG2B*, and *LRRC46*) are related to the immune system and *HEMK1* regulates methylation processes.

We detected 22 PSGs in wolverine, including six genes associated with energy production and conversion. Among them, *ATP6V0B, KMO*, and *SLC16A4* are primarily involved in insulin level regulation, and the metabolism of carbohydrates and fatty acids. Three PSGs (*OIP5*, *ZADH2* and *MTPAP*) are specifically associated with adipose tissue formation and intramuscular fat deposition. Additionally, we found three PSGs (*NBR1*, *TMEM38B*, *PPP1R18*) involved in selective autophagy as response to nutrient deprivation along with bone mass and density regulation, and resorption. We also detected PSGs (*DAB1*, *OPA1* and *CTNS*) linked to cognition, brain development and vision. Several PSGs (*BNIPL*, *IL18BP*, *CRNN*) were associated with the immune system; while three others (*ANAPC7*, *RNF212B*, *IZUMO3*) are involved in reproduction processes and *USB1* and *CLCN4* have a role in basal cell cycle processes. For the remaining two, *CEP95* and *FAM185A*, it was not possible to associate a specific phenotypic trait.

Among the 18 PSGs detected in the sable, three (*PRRT2*, *ATL2*, *SELENOI*) are associated with locomotion, and coordination, and *USP53* is associated with sensory perception and nervous system. Two PSGs, *VEGFC*, *RASA1*, are associated with blood vessel formation, three (*TTC4*, *ZBP1*, *CD247*) with the immune system and three (*IQUB*, *UBQLNL*, *MEIKIN*) with reproduction. Several PSGs (*EEF2KMT*, *DEUP1*, *ECD, IQCK*) are associated with cell cycle processes, and *ZC2HC1C* and *CCDC17* could not be associated with a particular biological process.

### Gene family expansions and contractions

Adaptive divergence between species may also be caused by changes in gene family sizes that occur during genome evolution and are due to gains (expansions) or losses (contractions) of genes or groups of genes (Olson, 1999; Tigano et al., 2020). All species analysed displayed more gene family contractions than expansions, with the wolverine having the highest contraction rate. This is most likely an artefact resulting from the fragmented genome assembly of this species (SI Figure S7). All identified expansions were in the form of gene duplications, with one putative triplication detected in tayra (SI Table S5B).

Tayra and sable had similar numbers of gene family expansions and contractions (SI Figure S7): 34 expansions and 169 contractions in tayra, and 33 expansions and 162 contractions in sable. The less contiguous wolverine genome contained 7 expansions and 649 contractions (SI Tables S5A, S5B). Due to the stochastic nature of gene losses and the potential inflation of estimates resulting from different genome assembly contiguities, we here focus on gains of gene copies.

Expanded gene families in the tayra genome are associated with reproduction, metabolism, nervous and immune system, and cell cycle, among others (Figure 4C-D; gene names, descriptions and functions are given in SI Table S5B). Of the three reproduction-related genes, *SLC38A2* regulates supply of nutrients for fetal growth through the placenta during the peri-implantation period. The second, *HSD17B10*, is associated with regulation of pregnancy-sustaining steroid hormones, and *RBP2* is involved in retinol binding and vitamin A metabolism, necessary for oogenesis and embryogenesis, as well as vision. Four genes (*ATP6V1D*, *DBX2*, *SLC38A1, MAPKAPK5*) are associated with cerebral cortex development, synapse formation, visual perception, and learning processes. *ANKRD13A* is also associated with vision, more specifically with lens fiber generation and vitamin A metabolism. *N6AMT1* is involved in modulation of arsenic-induced toxicity. The olfactory receptor gene *TAAR5* is involved in behavioral responses in mammals, and was duplicated in both tayra and sable. We detected one putative triplication of *FKBP3*, a gene associated with immunoregulation, predominantly of T-cell proliferation. *PRR11* could not be associated with a particular biological process.

In the wolverine, two duplicated genes are related to the nervous system: *GFRA4* is implicated in motor neuron development and *KCNS1* in regulating mechanical and thermal pain sensitivity. *MTM1* is associated with positive regulation of skeletal muscle tissue growth and *MON1B* is implicated in immune response to viral infection.

In the sable, expansions involve gene families associated with the nervous system, specifically with sensory perception and locomotion, with angiogenesis, hair follicle development, and the immune system. *DUSP8* is involved in neuronal development, vision, olfaction, and spatial memory formation. *FBXL3* is associated with regulation of the circadian clock and *TIMM10* with hearing. Two genes, *CDC42* and *TCHHL1*, are implicated in hair-follicle development. Additionally, *CDC42*, a gene coding for a cell division control protein, is also involved in angiogenesis and hematopoiesis, alongside *TNFRSF12A* and *LXN*.

### Structural variation

Structural variants (SVs) modify the structure of chromosomes and can affect gene synteny, repertoire, copy number and/or composition (e.g. gain or loss of exons), create linkage-blocks, and modify gene expression (Chiang et al., 2017; Mérot et al., 2020), leading to complex variation in phenotypes and genetic diseases (Weischenfeldt et al., 2013). We investigated four types of SVs (deletions, duplications, insertions, inversions) in the three Guloninae.

We identified the highest number of species-specific SVs in sable (22979), followed by tayra (8907), and wolverine (264) (Figure 5A). The most abundant SVs detected in all three species are deletions (> 50bp), ranging from 183 species-specific deletions in wolverine to 21713 in sable. Duplications were the least frequent SV type among the three species (SI Figure S8A). For all three species, the majority of SVs are located in intergenic regions (> 80%), with a smaller portion found in genic regions, completely or partially overlapping protein-coding genes (UTRs, exons, introns). According to Variant Effect Predictor (VEP) classification, SVs impacting genic regions are classified either as high-impact variants or modifiers (McLaren et al., 2016) with putative consequences on gene transcription ranging from transcript truncation to transcript ablation or amplification. The highest number of species-specific genic SVs was detected in tayra, with 330 (3.70% of species-specific SVs), followed by 156 (0.68%) in sable and 53 (20.08%) in wolverine (SI Figure S8B). Other than the well documented impact of inversions on intra and interspecific gene flow (Porubsky et al., 2020; Wellenreuther & Bernatchez, 2018), determining the impact of inversions overlapping large sets of genes is still challenging, as the largest effect is likely to be restricted to genes near SV breakpoints. Therefore, we restricted our examination of gene function to loci affected by deletions, duplications, and insertions (Figure 5B; gene names, descriptions and functions are given in SI Table S6).

**Figure 5.**
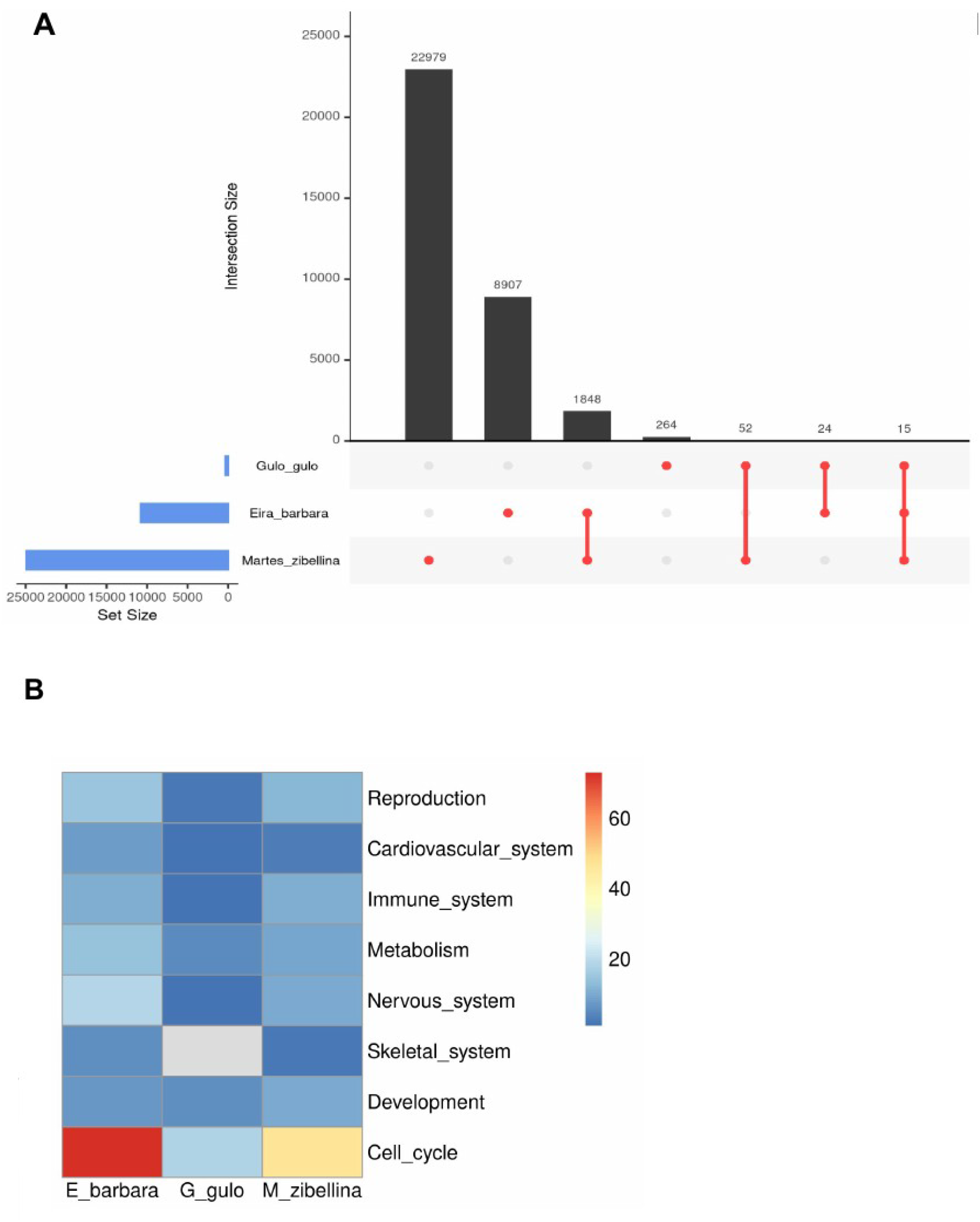
Structural variants detected in gulonine species. A) Shared and species-specific structural variants detected in wolverine (*Gulo gulo*), tayra (*Eira barbara*), and sable (*Martes zibellina*), B) Functional groups of genes affected by species-specific structural variants in 3 gulonine species (SV types: DEL, DUP, INS). Heatmap scale represents the number of genes. Heatmap cells in grey color indicate no observations for a given variable.

In the tayra genome, we observed 14 duplications spanning a combined length of 2.92 Mb, putatively affecting 24 protein-coding genes. Duplicated genes and gene blocks are associated with reproduction, olfaction, metabolism and energy conversion. This included *RNASEH2B*, a gene involved in *in utero* embryo development, and two genes involved in spermatogenesis, *DIAPH3* and *PCNX1*, with the latter an example of a complex SV involving heterozygous duplication and deletion of an exon (SV ~2kb in length). We detected 212 deletions in the tayra genome in relation to the domestic ferret reference, comprising a total length of 2.08 Mb, and affecting 247 genes, which are associated with reproduction, metabolism/energy conversion, nervous system, and cell cycle processes, among other functional categories (SI Table S6). Genes involved in placenta development and *in utero* embryogenesis include *HSF1*, *RSPO2*, and *DNMT3A*. Additionally, we detected *NLRP1* and *NLRP8*, both associated with preimplantation development, and highly expressed in oocytes. One short insertion was observed in *LIX1L*, a gene associated with anatomical structure morphogenesis.

In the wolverine genome, no duplications overlapped genic regions. However, 47 deletions spanning a combined length of 229 kb are putatively associated with transcript truncation or ablation in 48 genes. The majority of affected genes are associated with metabolism/energy conversion, development and basic cell cycle processes. This includes *GLUD1*, a gene involved in amino-acid induced insulin secretion, also found to be affected by a shorter deletion in sable, and *NSDHL*, a gene regulating cholesterol biosynthesis. Additionally, we detected deletions affecting *PARVA*, a gene associated with angiogenesis and smooth muscle cell chemotaxis and *DNAJC7*, involved in positive regulation of ATPase activity and regulation of cellular response to heat. We also detected one insertion in a gene of unknown function.

In the sable genome, we detected eleven duplications spanning a combined length of 324 kb, overlapping 16 genes associated with sensory perception, development, cell cycle, and the immune system. The 130 detected deletions (combined length of 408 kb) overlap 125 protein-coding genes associated with reproduction, immune system, development, metabolism, sensory perception, and cell cycle. Deletions were identified in two genes involved in keratinocyte differentiation, *PPHLN1*, also affected in wolverine, and *IVL*, associated with hair follicle development. Additionally, two short insertions were found in *NCOA4*, and *YIPF5*, genes associated with mitochondrial iron homeostasis and protein transport, respectively.

## Discussion

Here, we present a highly contiguous genome for the tayra (*Eira barbara*). Contiguity of the assembly and its gene completeness are similar to or higher than those of other carnivoran species using the same sequencing approach (Armstrong et al., 2019, Etherington et al., 2020, Kim et al., 2020), confirming the utility of linked-reads for assembly of mammal genomes.

Phylogenomic relationships among the mustelids resulted in a tree topology and divergence time estimates in agreement with previous studies using fewer loci (Koepfli et al., 2008; 2018; Li et al., 2014; Sato et al., 2012). We estimated the split between *Mustela* and Guloninae occurred 11.2 million years ago (Mya) (HPDI 13.1 - 9.5 Mya), followed by the split between *Eira* and the *Gulo-Martes* group 7.5 Mya (HPDI 9 - 6.1 Mya), and the split between *Gulo* and *Martes* at 5.9 Mya (HPDI 7.4 - 4.7 Mya).

Contrary to the findings by (Weissensteiner et al., 2020) in corvids, we did not observe a positive relationship between variation at the nucleotide level (heterozygous SNPs) and variation at the structural level (heterozygous SVs) within the gulonine species. The tayra displayed the highest nucleotide diversity (1.89 SNPs per kbp), but only the second highest amount of heterozygous SVs (2543, 23.6% of the total SVs, SI Figure S9). The sable had the second highest nucleotide diversity (1.44 SNPs per kbp), but the highest number of heterozygous SVs (14823, 59.5% of total). The wolverine displayed the lowest variation for both (0.28 SNPs per kbp, and 153 or 43.1% heterozygous SVs in total). It is known that SV calling using short-read data can miss a large number of SVs (Ebert et al. 2021). The fact that we did not detect a positive correlation between variation at the nucleotide and structural level, as would be expected if diversity of SNPs and SVs are correlated with population size, may result from our SV analysis relying on short-read data only. Weissensteiner et al. (2020), who did report a positive correlation between SNP and SV diversity, performed long-read based SV typing.

Assessment of variation among genome assemblies of closely related species also strongly relies on the contiguity and completeness of analysed assemblies (Gurevich et al., 2013; Totikov et al., 2021). This needs to be accounted for when examining variation among discontiguous genome assemblies. Here, the low contiguity of the wolverine assembly has likely impacted the number of PSGs and gene family expansions/contractions detected. Multiple, short insert size libraries sequenced to low coverage have likely resulted in decreased SV detectability. However, low diversity in the Scandinavian wolverine population is expected given previous results (Ekblom et al. 2018) and the species’ historical demography.

### Adaptive genomic variation

Among positively selected genes, gene family expansions and coding regions impacted by SVs, we found numerous candidate loci related to species-specific traits in Guloninae.

The three gulonine species differed with respect to which reproduction-related processes were associated with candidate genes. In the aseasonally breeding tayra, genes detected in three analyses (PSGs, gene family expansions, SVs) were frequently pregnancy-related, while in the two seasonal breeders, sable and wolverine, genes were involved in a greater diversity of reproduction-related processes, primarily in gametogenesis (Figure 6A). This may reflect the transition from seasonal to aseasonal breeding in the tayra lineage, the only extant aseasonal breeder among the Guloninae.

**Figure 6.**
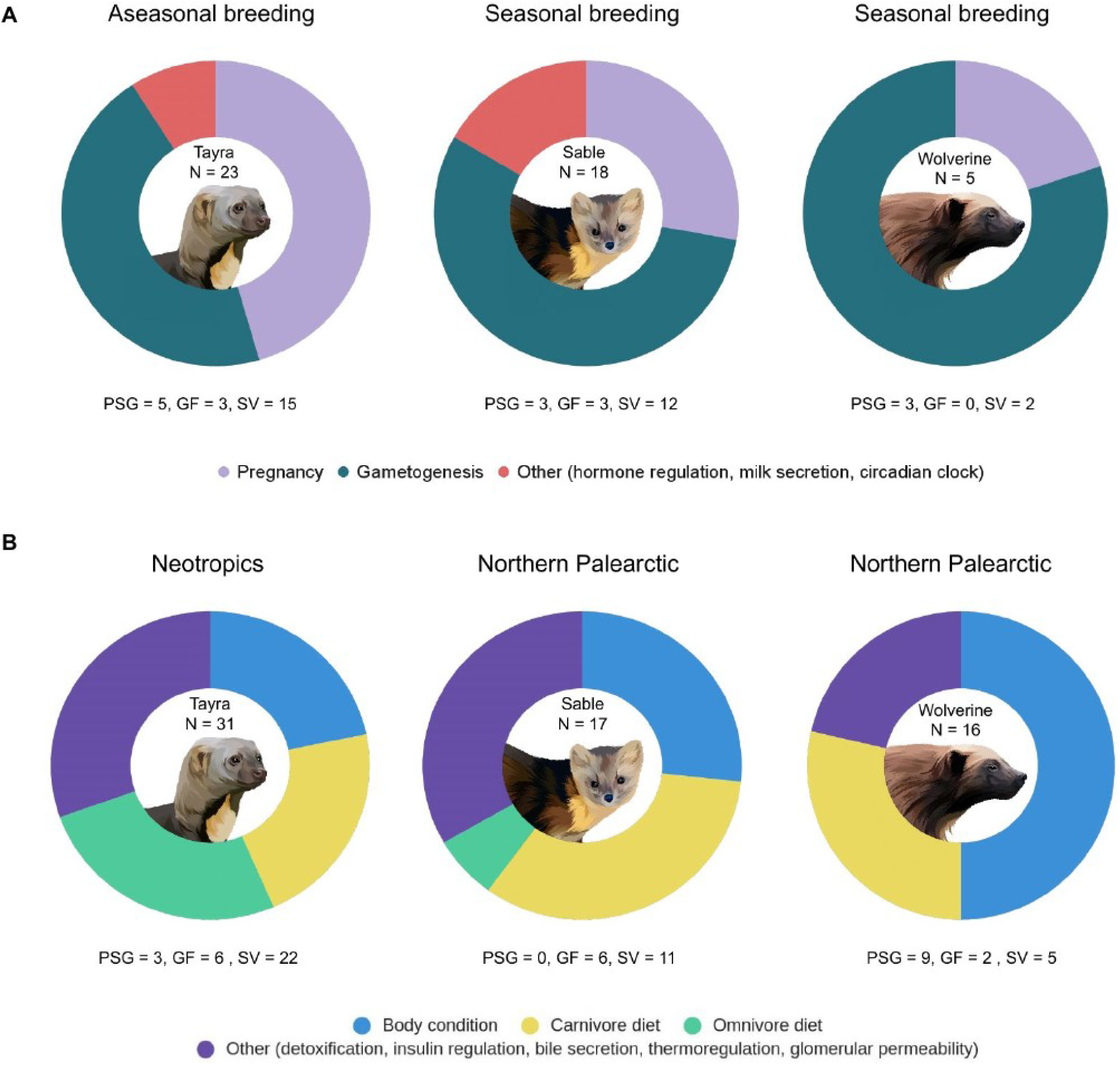
Summary of functional categories of (A) reproduction and (B) metabolism-related genes derived from analyses of positively selected genes (PSG), gene family expansion (GF) and structural variation (SV). N represents the total number of detected genes. Vector graphics of species are created based on royalty-free images (source: Shutterstock).

Shifts in dietary preferences have been linked to positive selection in single genes (Kosiol et al., 2008) and to copy number variation in metabolism-related gene families in mammals (Hecker et al., 2019; Rinker et al., 2019). Among Guloninae, the dietary gradient from omnivory in the neotropics to (hyper)carnivory in the northern Palearctic is reflected in the proportion of metabolism-related candidate genes observed (Figure 6B). The highest proportion of genes involved in carbohydrate metabolism (“omnivorous diet”) were detected in tayra, followed by sable, and none detected in wolverine. However, the highest number of genes involved in maintaining a stable body condition through shifts in energy sources (autophagy, fat storage and bone resorption) was observed for the wolverine, potentially reflecting this species’ adaptive response to unfavourable environmental conditions in its circumpolar habitat.

We note that in two analyses (PSGs and gene family evolution), we only considered variation in single-copy orthologs, not the entire gene repertoire of these species. Thus, our results are likely only an incomplete reflection of the genes involved in these traits.

### Seasonal breeders in the north palaearctic: wolverine & sable

Obligate embryonic diapause or the delayed implantation of the blastocyst is a widespread reproductive strategy among mustelids and other carnivorans. The majority of mustelids in the Northern Hemisphere are seasonal breeders, which includes wolverines and sables that undergo a prolonged period of a delayed implantation lasting several months (Mead, 1981; Svishcheva & Kashtanov, 2011). There is a strong selection for short birthing periods at higher latitudes (Heldstab et al., 2018), because it is one of the most energetically demanding periods, and its timing in relation to prey abundance has a critical impact on reproductive fitness (Inman et al., 2012).

Candidate genes potentially involved in seasonal breeding in wolverine and sable were associated with pregnancy, gametogenesis and circadian rhythm (Figure 6A). These mustelids occupy large home ranges, display territoriality and low population density, leading to infrequent encounters between males and females (Inman et al., 2012; Kashtanov et al., 2015). Thus, a mechanism that facilitates induced ovulation during the encounter would be an adaptive advantage (Larivière & Ferguson, 2003). We detected signals of positive selection in *ANAPC7* in wolverine, a gene involved in progesterone-mediated oocyte maturation and release from cell arrest prior to fertilization (Papin et al., 2004; Reis et al., 2006), that may have a role in increasing progesterone secretion and renewed embryonic development, as observed in skunks and mink (Mead, 1989).

Delayed implantation serves as an important adaptation to ensure maximal survival chances of offspring depending on the environmental conditions (Heldstab et al., 2018). Previous studies in mink (Lopes et al., 2003) showed that increased levels of vascular endothelial growth factors (VEGF) and their receptors correlate with the implantation process. *VEGFC*, primarily associated with angiogenesis and regulation of permeability of blood vessels during embryogenesis, was positively selected in sable, suggesting its possible involvement in embryo implantation regulation in this species.

Changes in testicular activity and spermatogenesis strongly correlate with season in mink (Blottner et al., 2006), and lynx (Jewgenow et al., 2006). We observed spermatogenesis-related genes affected by different types of variation in the three gulonine species. In the wolverine, positively selected genes involved in spermatogenesis included *IZUMO3* and *RNF212B*. Testes-expressed *IZUMO3* is essential for gamete fusion during fertilization (Ellerman et al., 2009) and positive selection of *IZUMO3* has been linked to species-specific adaptations rather than sexual selection (Grayson & Civetta, 2012) *RNF212B* is critical for successful crossing over in gametes, and knockout of *RNF212* causes sterility in mice (Reynolds et al., 2013). In sable, candidate genes involved in spermatogenesis were *UBQLNL* (Marín, 2014) and *SEPT12*, with the latter also being duplicated. Furthermore, *MEIKIN* and *CDK2*, both involved in meiosis, show signals of positive selection and rapid evolution through gene family expansion, respectively. *MEIKIN* has been found to be consistently rapidly evolving in other eutherian mammals (Pontremoli et al., 2021).

Seasonal breeding in many mammals is largely under photoperiod regulation, suggesting that the circadian system and pineal melatonin secretion play an important role in this reproductive strategy. Maintaining elevated melatonin in mink and spotted skunks has been shown to prolong embryonic diapause (Murphy & Fenelon, 2020). *FBXL3*, associated with maintenance of circadian clock oscillation in mammals (Fenelon et al., 2017; Shi et al., 2013) is duplicated in sable, indicating its possible relation to diapause regulation.

### Aseasonal breeder in the neotropic: tayra

In tropical regions, reproductive seasonality is less pronounced as environmental conditions are relatively stable throughout the year (McNutt et al., 2019). Tayras are aseasonal breeders with multiple estrous cycles per year (Proulx & Aubry, 2017) and do not exhibit embryonic diapause (Poglayen-Neuwall et al., 1989). The tayra represents the most basal taxon of the Guloninae, and is an exception regarding its reproductive strategy.

Genes that may be involved in the aseasonal breeding of tayra include *ETV2*, *MUC15*, *SLC38A2*, *HSD17B10*, *RBP2* and *RNASEH2B*. *ETV2* and *MUC15* are involved in placental and embryo development, and regulation of implantation (Shyu et al., 2007; Singh et al., 2019). The *MUC15* protein has been associated with an anti-adhesive function, and prevention of premature blastocyst attachment in rats (Denker, 2000; Poon et al., 2014). Increased expression of the neutral amino acid transporter *SLC38A2* has been detected in the endometrium of cattle during the preimplantation period (Forde et al., 2014), and in mice placentas during late gestation, maintaining fetal growth when maternal growth is restricted by undernutrition (Coan et al., 2010). *SLC38A2* is a duplicated and rapidly evolving gene in tayra, suggesting a potential role in supporting multiple pregnancies during the year. Duplication of *HSD17B10* might indicate a gene dosage modulation in steroid hormone-regulated pregnancy in tayra. This gene is involved in mitochondrial fatty acid metabolism, cell growth and stress response (Liu et al., 2020). It is highly expressed in fetal and maternal liver, influencing the maintenance of human pregnancy and providing protection against excitotoxicity (Hill et al., 2011). Besides being associated with vision, retinol binding protein *RBP2* has a significant role in regulating retinoids during oogenesis and embryogenesis, and positively impacts oocyte maturation in mice, cattle, pigs and sheep, (Brown et al., 2003; Harney et al., 1993). It was found to be duplicated in tayra, as well as a partial duplication detected in *RNASEH2B*. Mutations in this gene have been shown to cause interferonopathies, leading to impaired fetal brain development (Crow et al., 2015; Yockey & Iwasaki, 2018). In tayra, duplication of this gene potentially increases gene expression and may assist with maintenance of embryo development. Furthermore, five genes involved in placental development, implantation and embryogenesis (*HSF1*, *RSPO2*, *NLRP1*, *NLRP8* and *DNMT3A*) have been affected by partial deletions in tayra, suggesting further changes in dosage of pregnancy-related genes.

### Resource availability in the northern Palaearctic: wolverine & sable

Overcoming demanding periods and staying active throughout the year is challenging for non-hibernating northern palearctic species and requires specific mechanisms to cope with seasonal food scarcity. Wolverines have evolved several adaptations to efficiently exploit scarce food resources and overcome constraints of the low-productivity environment (Inman et al., 2012).A critical metric of body condition is the amount of stored fat, representing an important energy source between feeding opportunities and providing thermal insulation in cold environments. Undernutrition detrimentally affects reproduction, and may lead to decreased survival rates (Brown & Lasiewski, 1972; Schulte-Hostedde et al., 2005). Three PSGs detected in the wolverine are involved in adipose tissue formation, *MTPAP, OIP5 and ZADH2* (Han et al., 2012; Inoue et al., 2014; Yu et al., 2013), and may be associated with selective fatty acid mobilization stimulated by fasting periods (Inman et al., 2012; Krebs et al., 2004), as observed in mink (Nieminen et al., 2006), and raccoon dog (Mustonen et al., 2007).

One of the responses to prolonged periods of nutrient deprivation and extreme environmental conditions is suppressed bone resorption and formation (Lennox & Goodship, 2008; McGee-Lawrence et al., 2015). Beside regulating survival of blastocysts during delayed implantation (Lee et al., 2011; Lim & Song, 2014), it has been demonstrated that autophagy plays a critical role in maintenance of bone homeostasis (DeSelm et al., 2011; Montaseri et al., 2020). It is thus of note that genes involved in bone mass regulation, resorption (*PPP1R18*, *TMEM38B*) and autophagy (*NBR1*) are under positive selection in wolverines. We also detected a duplication of the muscle growth regulating gene *MTM1*. While a lack of *MTM1* will lead to muscle hypotrophy through unbalanced autophagy in humans and mice (Al-Qusairi et al., 2013), a gene duplication may facilitate muscle growth.

In sable, fatty acids are uniformly mobilized from anatomically distinct fat deposits (Nieminen & Mustonen, 2007), and we observed duplications of *ASB6*, *SLC25A10*, *RNF186* and *BORCS6*, which have been implicated in effective regulation of fat storage and response to nutrient availability (Mizuarai et al., 2005; Okamoto et al., 2020; Schweitzer et al., 2015; Wilcox et al., 2004). Moreover, partial deletions of *APOD*, *PDHB*, *LDLR*, and *CERS5*, associated with lipid metabolic processes (Carmo et al., 2009; Gosejacob et al., 2016; Serão et al., 2011; Tavori et al., 2015), indicate a possible association with adaptive fasting and energy conservation in this species.

We observed partial deletions in *DNAJC7*, a gene involved in thermoregulation (Sonna et al., 2002) in both sable and wolverine, but not in tayra, indicating a potential role in gulonines inhabiting colder environments. Furthermore, we detected partial deletions in *GLUD1* in sable and wolverine. *GLUD1* regulates insulin secretion (Fahien & Macdonald, 2011) and might play a role in mobilizing lipids, causing states of reversible insulin resistance as an adaptive response during prolonged periods of nutrient deprivation, as has been observed in some carnivorans (Martinez & Ortiz, 2017; Viscarra et al., 2013).

In sable and tayra, we observed a partial deletion of *PPP1CC*, associated with glycogen metabolism (Li et al., 2021), tentatively facilitating a more ‘omnivorous diet’ in these species compared to hypercarnivory in wolverines (Lopes-Marques et al., 2020; Sharma et al., 2018).

There are notable differences in hair follicle structure and fur density between palearctic and neotropic species (Kitchener et al., 2018), and we observed two gene copy gains in sable, *TCHHL1*, involved in hair morphogenesis (Wu et al., 2011), and *CDC42*, required for differentiation of hair follicle progenitor cells (Wu et al., 2006).

### Resource availability in the Neotropics: tayra

Tayras exploit diverse food sources and experience relatively stable resource availability all year round, as (sub)tropical habitats have higher primary productivity compared to habitats at higher latitudes (Heldstab et al., 2018; Zhou et al., 2011). We found candidate genes associated with regulation of fructose metabolism, consistent with dietary preferences of tayra involving a variety of fruit and honey (Kratzer et al., 2014). The uricase (*UOX*) gene, which is involved in regulation of purine metabolism and conversion of fructose to fat (Johnson & Andrews, 2010) was under positive selection in tayra. Old World monkeys show low levels of uricase enzymatic activity, and uricase is pseudogenized in humans, great apes, and New World monkeys due to silencing mutations (Oda et al., 2002).

High rates of lineage-specific variation in gene family size, especially those families involved in immune response or detoxification of xenobiotic molecules (Thomas, 2007), are likely associated with environmental changes during speciation (Lynch & Conery, 2000; Zhang, 2003). We found a duplication of *N6AMT1*, a member of a gene family implicated in environmental stress response. *N6AMT1* is associated with conversion of the arsenic metabolite monomethylarsonous acid to the less toxic dimethylarsonic acid (Ren et al., 2011). Chronic arsenic exposure through water and soil, has been monitored in the neotropics, as it represents a severe public health issue, leading to various diseases (Zhang et al., 2015). This duplication may represent an adaptation of tayra to this xenobiotic compound.

Finally, we also found candidate genes associated with vision and cognition, which may be associated with the tayra’s primarily diurnal foraging activity. Tayras supposedly detect prey primarily by smell, as their eyesight has been described as being relatively poor (Galef et al., 1976; Wilson & Mittermeier, 2009). This is contradictory to the observed behaviour of caching of unripe but mature stages of both native and non-native fruit by tayras (Soley & Alvarado-Díaz, 2011). We detected variation in several genes implicated with lens fiber formation and retinal vascularization, including gene expansions of *ANKRD13A* (Avellino et al., 2013) and *RBP2* (D’Ambrosio et al., 2011), suggesting that tayras might not in fact exhibit poor eyesight. The caching behavior mentioned above indicates remarkable sensory perception (Presley, 2000), and recognition of different stages of fruit development, likely evolved as a response to avoid competition with other species during seasons when fruit becomes ripe (Soley & Alvarado-Díaz, 2011). We observed genes related to memory and learning such as *SLC38A1* (Qureshi et al., 2019) and *CRBN* (Higgins et al., 2010) that may have been involved in the evolution of caching behavior in tayra.

Mustelids are a remarkable example of adaptive radiation, and we show how, together with positively selected loci, changes in gene family size and structural variants have shaped genomes in this diverse taxonomic group. We demonstrated that the latter two sources of variation contribute significantly to the number of loci potentially involved in adaptive genomic evolution. These variant types are often not considered in comparative genomic studies of non-model species, even though they impact more nucleotides than SNPs.

## Supporting information

Supplemental_Information_Derezanin

SI_tables_S4-S6_Derezanin

## Acknowledgments

We thank Dr. Rafael from the Felidae Wildkatzen- und Artenschutzzentrum Barnim for kindly providing the tayra sample, and Michael Hofreiter from the Adaptive Genomics group (University of Potsdam) for assistance in generating the 10x Genomics linked-read library.

David Duchêne was funded by a Carlsbergfondet postdoctoral fellowship (grant number CF18-0223). Sergei Kliver, Andrey Tomarovsky and Azamat Totikov were funded by the Russian Foundation for Basic Research (grant number 20-04-00808). Azamat Totikov and Andrey Tomarovsky were additionally funded by JetBrains Research.

## Data Accessibility and Benefit-Sharing Statement

The tayra primary and alternate genome sequences have been deposited in GenBank under the accession JAHRIG000000000-JAHRIH000000000. All data generated or analysed during this study are included in this published article and its supplemental information files. Bioinformatic scripts will be publicly available in a Github repository specifically designated to this study, with a DOI assigned to it. Raw sequencing data will be deposited and available for download from the NCBI SRA repository.

## Author contributions

LD conceived the study, conducted genome assembly, gene family evolution and structural variation analysis, interpreted the data, and wrote the manuscript. AB and SJ conducted positive selection analysis, and interpreted the data. PD carried out demographic history reconstruction and data interpretation. DAD designed and performed phylogenomic analysis and molecular dating, and wrote the manuscript. JHG conducted repetitive elements analysis and interpreted the data. SK, AT and AT performed reference-based scaffolding, alignment to pseudochromosome assemblies, sex verification and nucleotide diversity assessment, interpreted the data and worte the manuscript. KPK advised comparative genomic analyses, and edited the manuscript. DM supervised and performed gene family evolution analysis, and advised other comparative genomic analyses. MP conducted whole genome sequencing and sequencing data interpretation. JF and DWF conceived the study, interpreted the data, wrote and edited the manuscript. All authors have reviewed and approved the manuscript.

