## Supplemental_Information_Derezanin for "Multiple types of genomic variation contribute to adaptive traits in the mustelid subfamily Guloninae"

MOLECULAR ECOLOGY

### Table of contents:

**SUPPLEMENTAL FIGURES AND TABLES** **1**

Figure S1. Genome alignment between domestic ferret and tayra assemblies. 1

Table S1. Tayra genome assembly metrics. 2

Figure S2. Genome completeness metrics. 3

Table S2. Major types of transposable elements found in the genomes of four mustelid species. 4

Figure S3. Repeat landscape of the tayra genome assembly. 4

Figure S4. Historical demography of three gulonine species. 5

Figure S5. Coverage plots for (A) the tayra, (B) the sable, and (C) the wolverine. 6

Table S3. Statistics for counts of heterozygous SNPs. 7

Figure S6. Genome-wide heterozygosity for tayra (Eira barbara), sable (Martes zibellina) and wolverine (Gulo gulo). 8

Figure S7. Gene family expansions and contractions for eight carnivoran species. 8

Figure S8. Structural variants detected in gulonine species. 9

Figure S9. Heterozygous and homozygous SVs detected in each gulonine species. 10

**SUPPLEMENTAL INFORMATION** **11**

Genome assembly and processing of sequencing data 11

Alignment to pseudochromosome assemblies and sex verification 11

Phylogenomic data preparation, analysis and dating 11

Nucleotide diversity 13

Structural variants 14

**REFERENCES** **15**

**Supplemental Tables S4-S6 - References** **18**

SUPPLEMENTAL FIGURES AND TABLES

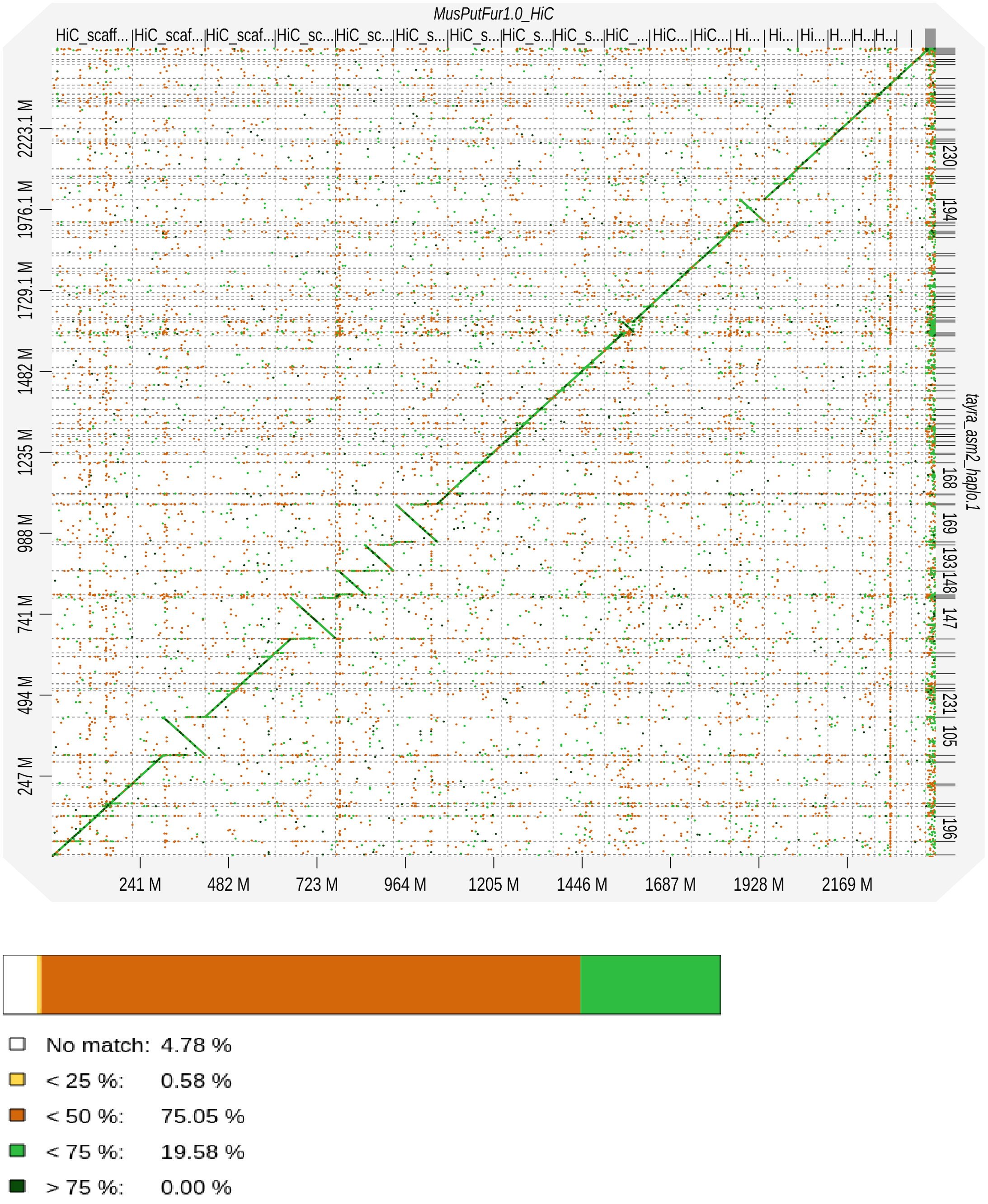


**Figure S1.** Genome alignment between domestic ferret and tayra assemblies.
Synteny analysis of domestic ferret Hi-C genome assembly (upper X-axis) and tayra assembly (right Y-axis) with overall match indicating 95% identity (with > 50% similarity).

**Table S1.** Tayra genome assembly metrics.
Combined assembly metrics obtained from Supernova assembler and Quality Assessment Tool for Genome Assemblies (QUAST).

| Supernova assembly metrics (v2.1.1) | |
|---|---|
| Sequencing strategy | Illumina NovaSeq + 10x Genomics |
| Assembly method | Supernova assembler v2.1.1 |
| Sequencing reads | 1.3 billion reads, 150 PE |
| Mean read length | 139.5 bases |
| Raw coverage | 75.63x |
| Effective coverage | 55.34x |
| Proper read pairs | 88.78 % |
| Median insert size | 344 bases |
| GC content | 41.76 % |
| Molecule length | 50.75 kb |
| Contig N50 | 289.96 kb |
| Scaffold N50 | 42.07 Mb |
| Number of scaffolds | 14579 |
| Number of scaffolds ($\geq$ 5 kb) | 3773 |
| Longest scaffold | 123.17 Mb |
| Phaseblock N50 | 5.77 Mb |
| Assembly size | 2.447 Gb (scaffolds $\geq$ 5 kb) |
| QUAST assembly metrics (v5.0.2) | |
| Scaffolds > 10 kb | 1621 |
| Scaffolds > 25 kb | 469 |
| Scaffolds > 50 kb | 254 |
| Scaffold L50 | 17 |
| N's per 100 kbp | 1145.00 |

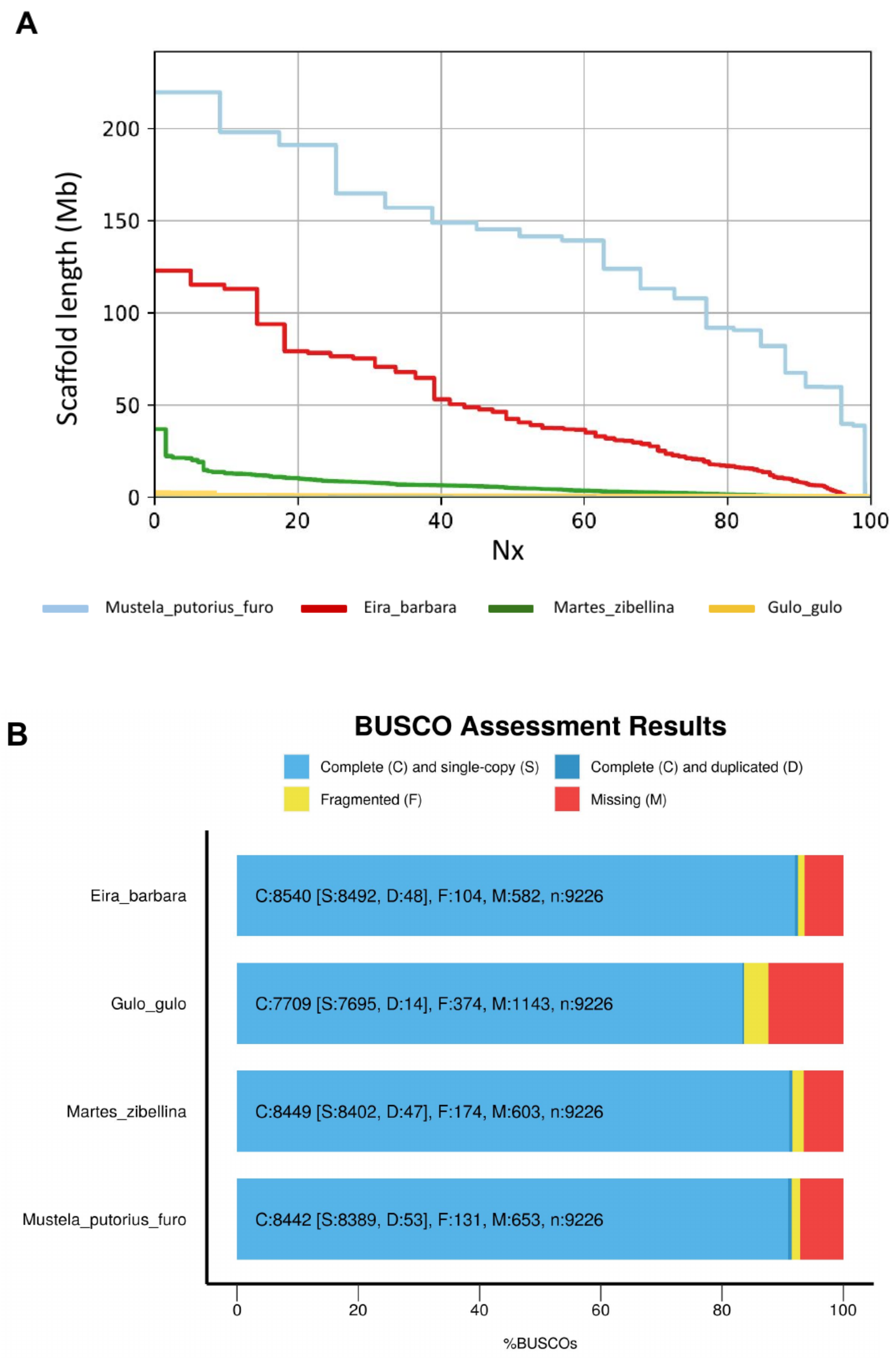


**Figure S2.** Genome completeness metrics.
A) Summary of QUAST assembly completeness analysis (number of contigs is represented on the X-axis) and (B) comparison of the BUSCO gene completeness assessment for four mustelid species.

**Table S2.** Major types of transposable elements found in the genomes of four mustelid species.

| Type | Tayra (*Eira barbara*) | Sable (*Martes zibellina*) | Wolverine (*Gulo gulo*) | Domestic ferret (*M. putorius furo*) |
|---|---|---|---|---|
| SINE | 1.91 % (47 Mb) | 1.5 % (36 Mb) | 1.56 % (37 Mb) | 1.36 % (32 Mb) |
| LINE | 23.17 % (570 Mb) | 19 % (465 Mb) | 15.73 % (381 Mb) | 17.13 % (417 Mb) |
| LTR | 4.05 % (99 Mb) | 3.5 % (84 Mb) | 3.28 % (79 Mb) | 4.4 % (106 Mb) |
| DNA | 1.63 % (40 Mb) | 2.03 % (49 Mb) | 2.05 % (49 Mb) | 3.13 % (75 Mb) |
| Total | 32.96 % (814 Mb) | 28.15 % (681 Mb) | 25.19 % (610 Mb) | 28.86 % (695 Mb) |

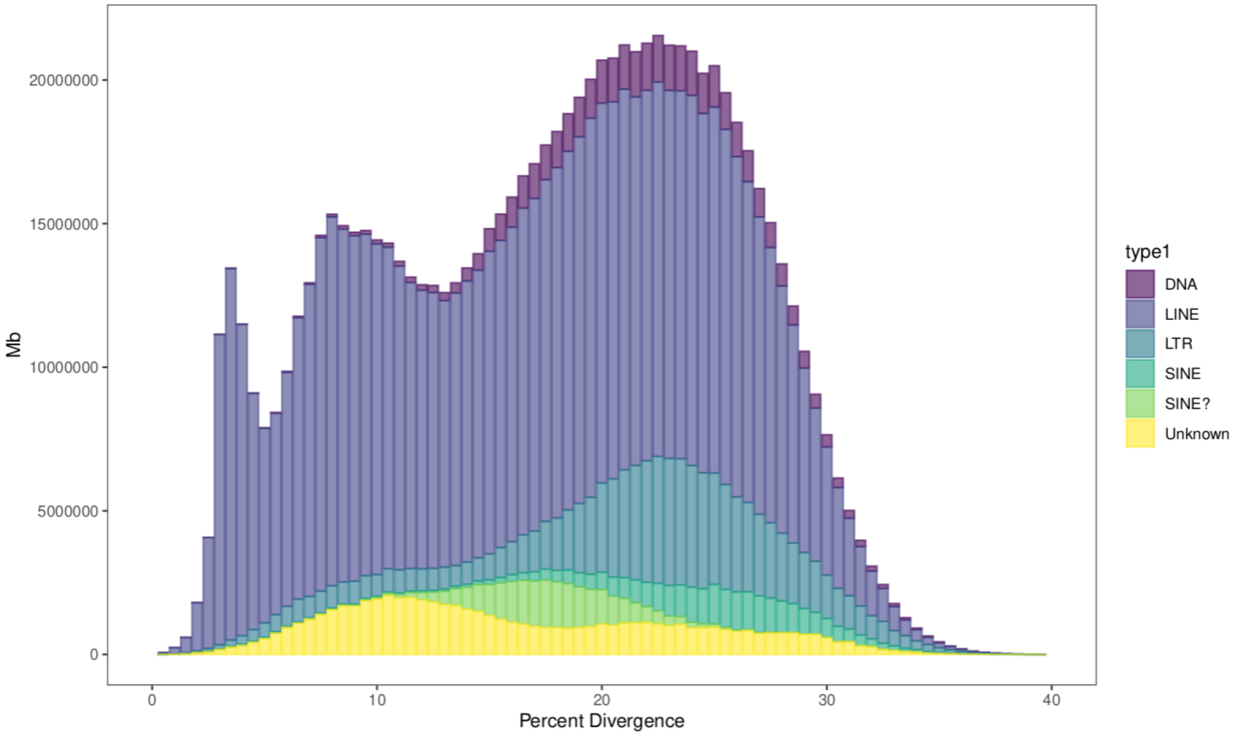


**Figure S3.** Repeat landscape of the tayra genome assembly.
Coloring scheme refers to different proportions of different repeat classes and x-axis indicates percent divergence within different repeat classes.

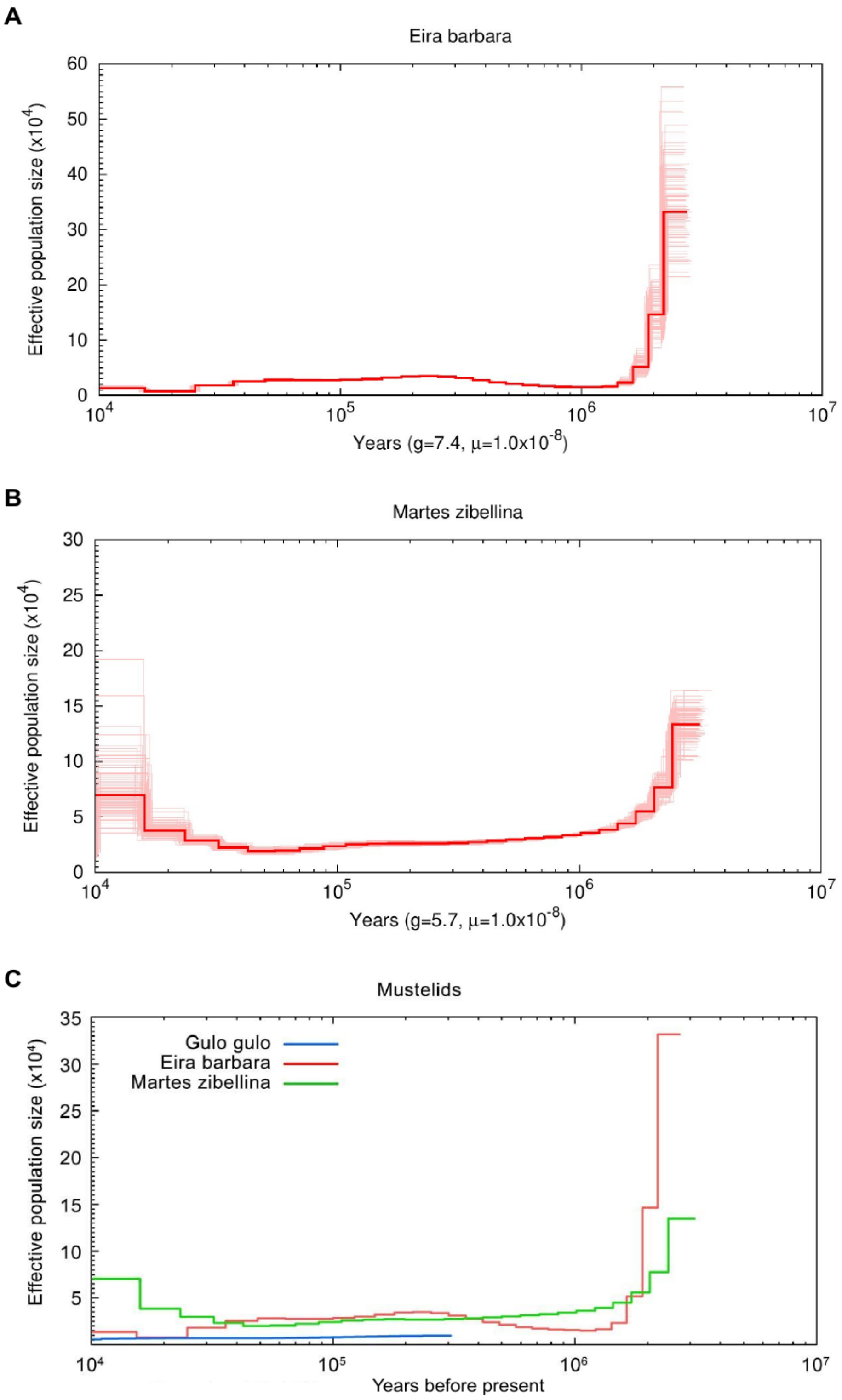


**Figure S4.** Historical demography of three gulonine species.

Bootstrapped inference of effective population size change over time for (A) tayra and (B) sable, and (C) historical demography inferred for all three species under different generation times (*G.gulo* = 6y, *E.barbara* = 7.4y, *M.zibellina* = 5.7y). Demographic trajectory for wolverine has been reconstructed from Ekblom et al. 2018. Time scale on the x-axis is calculated assuming a mutation rate of $1.0x10^{-8}$.

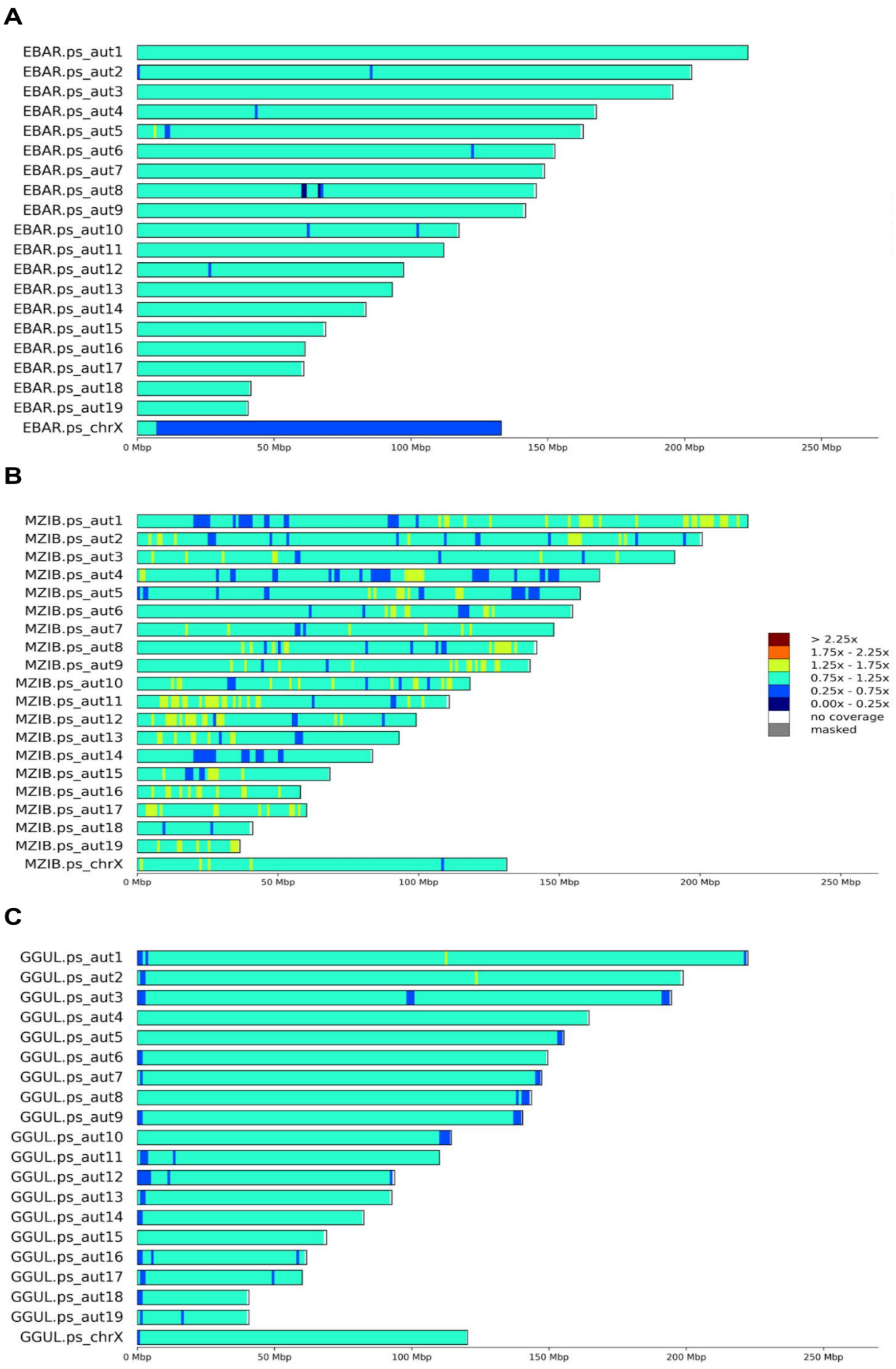


**Figure S5.** Coverage plots for (A) the tayra, (B) the sable, and (C) the wolverine.
Coverage was calculated in non-overlapping sliding windows of 1 Mbp and divided by whole genome median coverage.

**Table S3.** Statistics for counts of heterozygous SNPs.
SNPs are counted in 1 Mbp non-overlapping sliding windows for tayra, sable and wolverine scaled to SNPs per kbp. Bold marks the median values.

| Species | All scaffolds, heterozygous SNPs/kbp | | | | Without X pseudochromosome, heterozygous SNPs/kbp | | | |
|---|---|---|---|---|---|---|---|---|
| | Min | Max | Median | Mean | Min | Max | Median | Mean |
| *Eira barbara* | 0 | 5.99 | **1.89** | 1.81 | 0 | 5.99 | **1.93** | 1.90 |
| *Martes zibellina* | 0 | 4.45 | **1.44** | 1.51 | 0 | 4.45 | **1.47** | 1.56 |
| *Gulo gulo* | 0 | 1.24 | **0.28** | 0.27 | 0 | 1.24 | **0.29** | 0.28 |

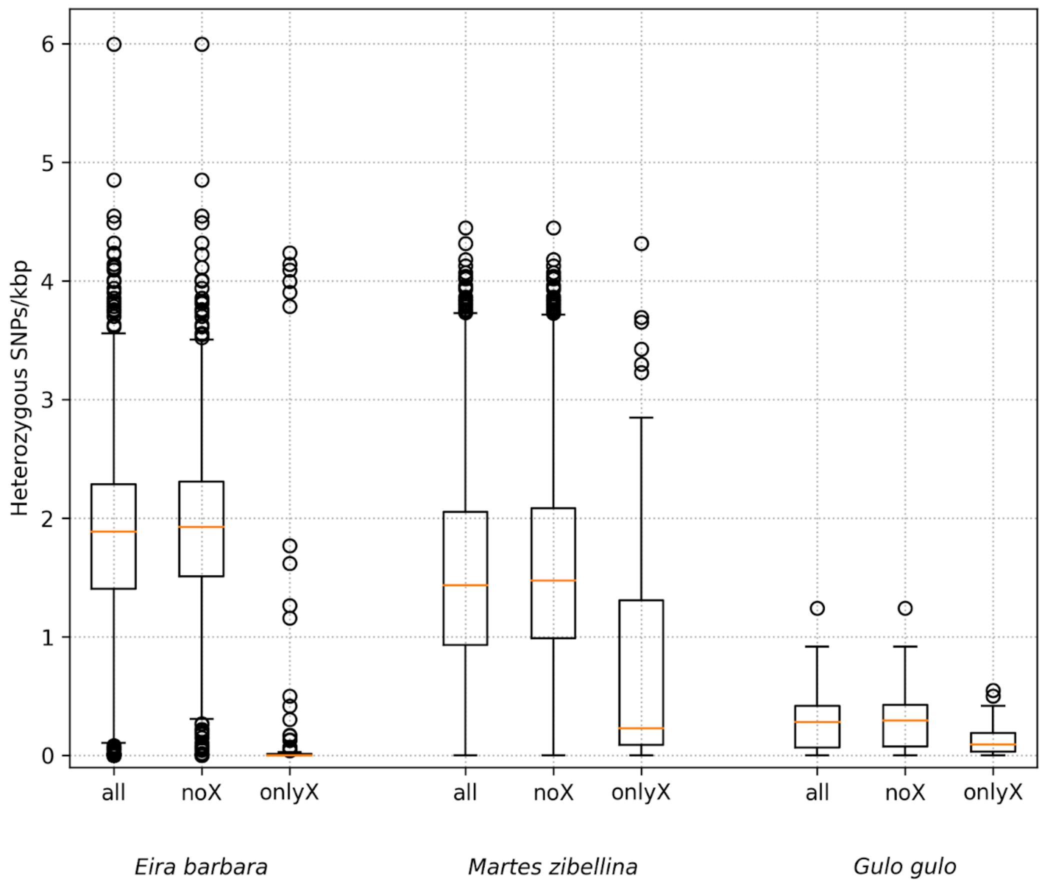


**Figure S6.** Genome-wide heterozygosity for tayra (*Eira barbara*), sable (*Martes zibellina*) and wolverine (*Gulo gulo*).
SNPs are counted in 1Mbp non-overlapping sliding windows and scaled to heterozygous SNPs per kbp. Corresponding pseudochromosome assemblies based on the domestic ferret assembly were used

as reference. 'all' includes windows from all scaffolds of at least 1 Mbp, 'noX' - without windows from the X pseudochromosome C-scaffold (ps_chrX), 'onlyX' - only windows from ps_chrX . Exclusion of ps_chrX slightly affected the borders of boxes and whiskers (all vs noX). Orange lines on boxplots correspond to median values, box edges - to 25th and 75th percentiles (Q1 and Q3), lower whisker - to max(0, Q1 - 1.5*IQR), upper whisker - to Q3 + 1.5*IQR, respectively. Interquartile range equals Q3 -Q1. Circles are outliers, i.e values outside [max(0, Q1 - 1.5*IQR), Q3 + 1.5*IQR ].

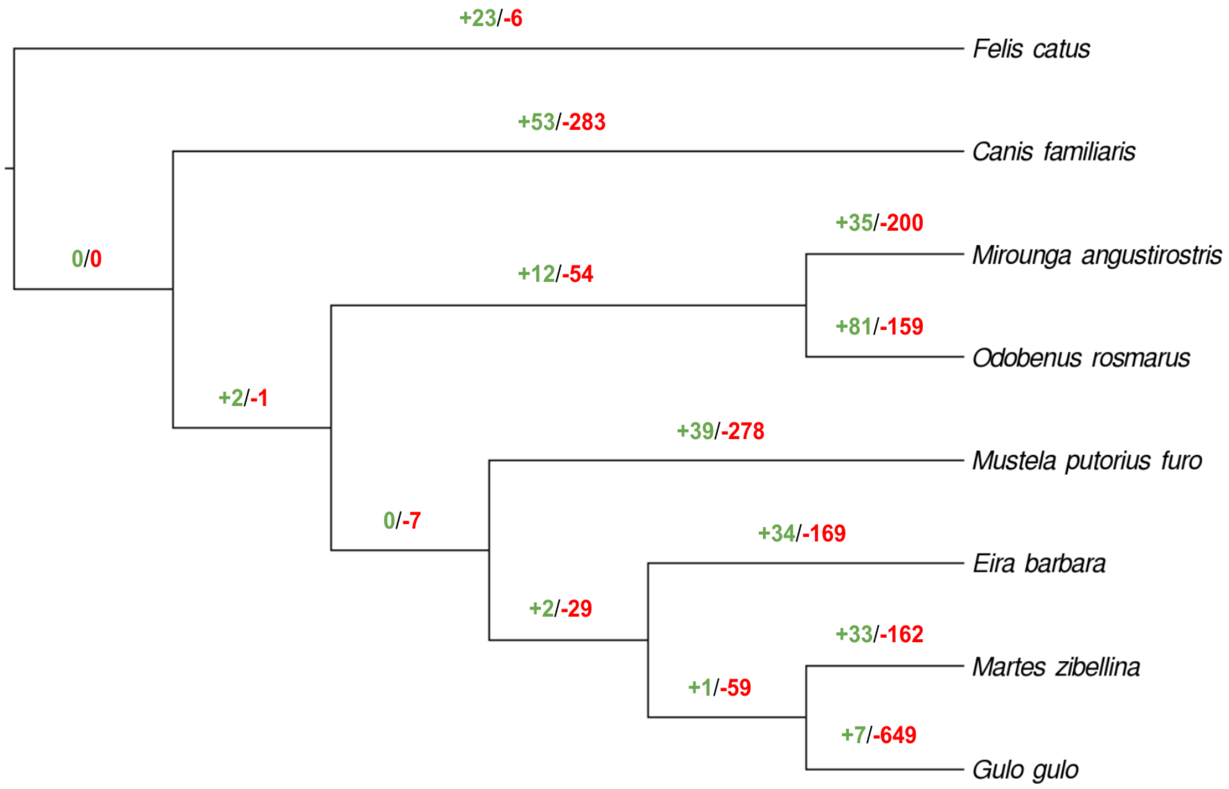


**Figure S7.** Gene family expansions and contractions for eight carnivoran species.
The species tree was built with FigTree v1.4.4 (https://github.com/rambaut/figtree). On each branch, the number of gene family expansions (+, green) and contractions (-, red) are represented. Counts are based on an error-corrected model of gene family evolution (analysed with CAFE).

A

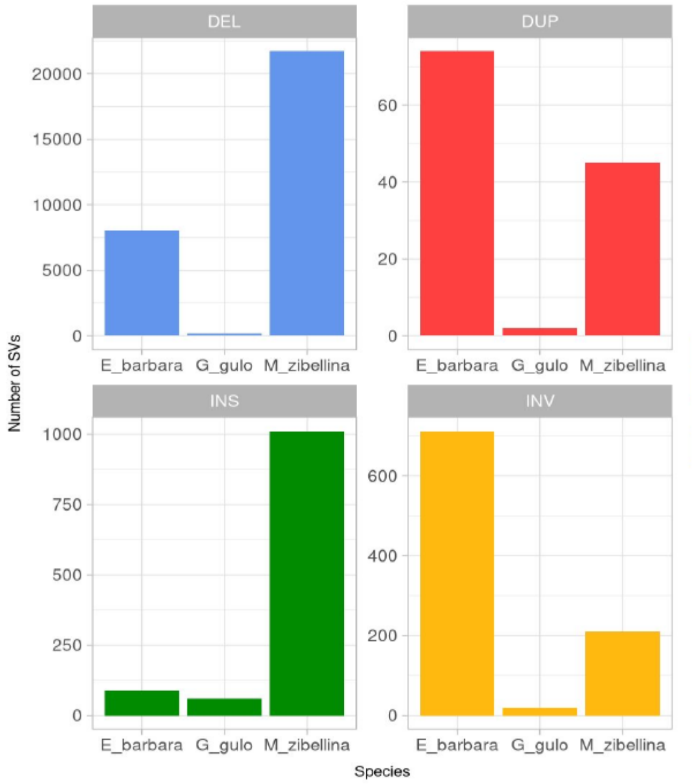


B

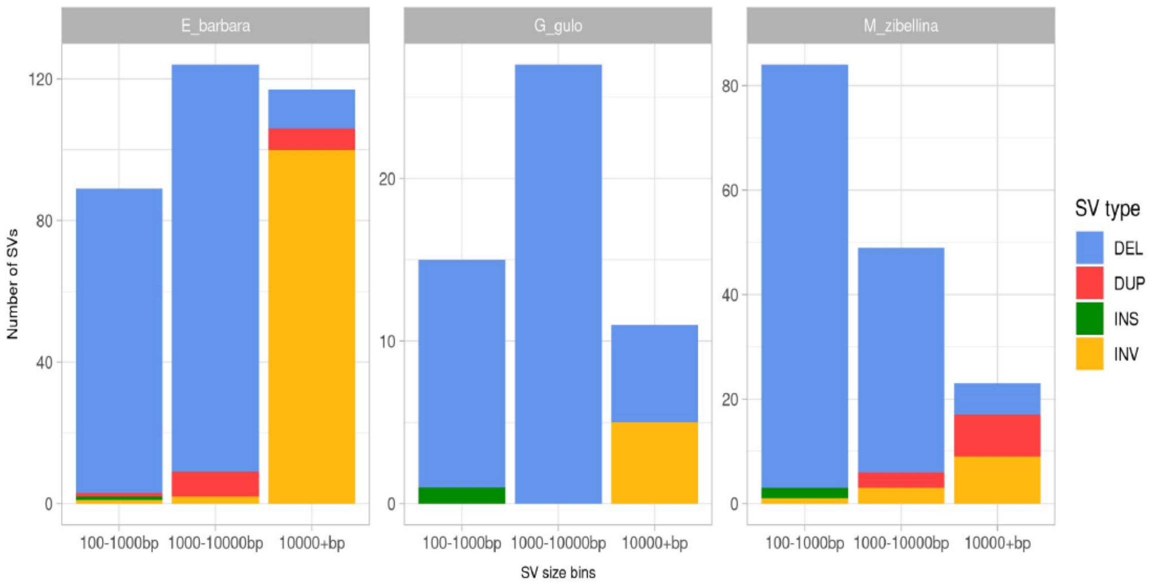


**Figure S8.** Structural variants detected in gulonine species.
**A)** Counts of different types of species-specific structural variants detected in tayra (E_barbara), wolverine (G_gulo), and sable (M_zibellina), **B)** Length distribution of species-specific structural variants overlapping genic regions detected in tayra (E_barbara), wolverine (G_gulo), and sable (M_zibellina).

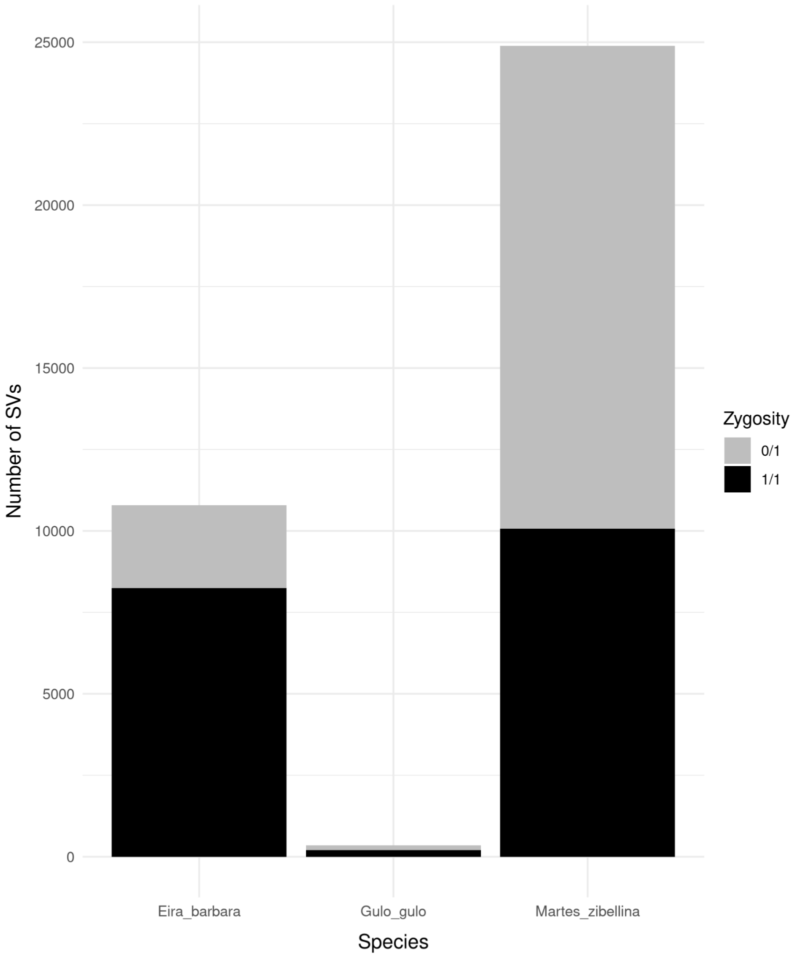


**Figure S9.** Heterozygous and homozygous SVs detected in each gulonine species. Heterozygous SVs represented with "0/1" (grey), homozygous SVs with "1/1" (black).

SUPPLEMENTAL INFORMATION

### *Genome assembly and processing of sequencing data*

Due to uncertainty regarding the optimum number of reads for genome assembly, we generated three *de novo* genome assemblies with default parameters, each with a different read input: 900 million, 1.3 billion and 1.7 billion paired reads, respectively. Based on Supernova assembly metrics (contig, scaffold, and phase block N50 sizes), we chose the assembly generated using 1.3 billion paired reads for further analysis (SI Table S1).

Prior to mapping, linked-read barcodes were trimmed from tayra sequencing reads with proc10xG (filter_Reads.py, https://github.com/ucdavis-bioinformatics/proc10xSC). For the other two species, we used the reads used for generating the respective genome assemblies of Scandinavian wolverine (*Gulo gulo*, GenBank ID: GCA_900006375.2, PRJEB10674), and sable (*Martes zibellina*, GCA_012583365.1, PRJNA495455). Adapter clipping and quality trimming (Q30, min. length 80 bp) were performed on all samples with TrimGalore v0.6.4 (Krueger et al., 2021). The proportion of reads mapping to the reference domestic ferret genome for all three samples was above 96%. Insert size distributions for each sample library were generated with Svtyper v0.7.1 (Chiang et al., 2015).

### *Alignment to pseudochromosome assemblies and sex verification*

Trimmed reads were aligned to the pseudochromosome assemblies of corresponding species using BWA version 0.7.17 (Li & Durbin, 2009). Read duplicates were marked using the markdup utility from Samtools version 1.10 (Li, 2011). To verify the sex of the tayra, sable and wolverine genomes, two approaches were applied: marker-based and coverage-based. For the marker approach, we used the Y-chromosome-specific SRY gene (sex determining region Y). The amino acid sequence of the *Martes melampus* SRY (GenBank ID: BAJ05096.1; Yamada & Masuda, 2010) was downloaded from the NCBI protein database and aligned to the pseudochromosome assemblies using Exonerate v 2.2.4 (Slater & Birney, 2005) in protein2genome mode to identify orthologous sequences. For the second approach, a per-base genome coverage was estimated using Bedtools v2.29 (Quinlan & Hall, 2010). Mean and median values for both non-overlapping sliding windows of 1Mbp and whole genomes were calculated and visualized using scripts from the MACE package (https://github.com/mahajrod/MACE).

### *Phylogenomic data preparation, analysis and dating*

We performed sequence alignment of 6020 coding genomic regions of single-copy orthologs shared across eight carnivoran taxa. Our taxon set included domestic cat (*Felis catus*), domestic dog (*Canis familiaris*), northern elephant seal (*Mirounga angustirostris*), and walrus (*Odobenus rosmarus*), in addition to our four mustelid species. Each genomic region was first filtered for highly divergent segments of sequences using the option trimNonHomologousFragments in MACSE v2 (Ranwez et al., 2011). This was followed by coding-region-aware sequence alignment using the alignSequences option. Sequences that were highly divergent and suspected of suffering from problematic alignment or sequencing were also identified using TreeShrink (Mai & Mirarab, 2018), based on a phylogenetic gene tree estimated using IQ-TREE v2 (Minh et al., 2020b). This was followed by a second round of alignment excluding the flagged sequences. We also minimized alignment error by excluding codons with >50% of missing data, and with heterozygosity in the translated amino-acid sequences above >50%.

Possible sources of biased phylogenetic inferences due to mis-specification of the substitution model were minimized by assessing model adequacy for each sequence alignment. We performed assessment of model adequacy using methods based on simulations (Duchêne et al., 2018a) and divergence matrices (Naser-Khdour et al., 2019) as implemented in PhyloMAd (Duchêne et al., 2018b) and IQ-TREE v2. Sequence alignments were retained if they passed all tests of substitution model adequacy and retained at least four taxa and 100 nucleotides. Gene trees were estimated from the acceptable gene regions by first selecting the best substitution model from the GTR+F+Γ+I+R family (Kalyaanamoorthy et al., 2017) and calculating approximate likelihood-ratio test (aLRT) branch supports (Anisimova & Gascuel, 2006), as implemented in IQ-TREE v2. Sequence alignment, cleaning and model adequacy assessment led to a phylogenomic data set with 2457 gene regions comprising over 3.2 million nucleotide sites.

Species tree estimates were performed using two methods. First, we concatenated sequence alignments and performed an analysis assuming that differences between gene trees are caused exclusively by stochastic error arising from having a finite sample of sites. Analysis of the concatenated data set was performed by taking the best model selected per gene region, and a model where branch lengths can vary among gene trees but maintain their relative lengths among branches (Duchêne et al., 2020). Second, we performed species tree inference under the multi-species coalescent using ASTRAL-III (Zhang et al., 2018), assuming that gene tree discordance arises from ancestral population-level processes. Gene trees were used as input for this analysis, collapsing into polytomies all aLRT branch supports below 50 prior to analysis to minimize the impact of stochastic error in gene trees. The estimate of the species tree was accompanied by local posterior probabilities as metrics of branch support.

Concordance factors were calculated using IQ-TREE v2 to explore the decisiveness of the phylogenomic signals found across gene trees (gCF) and alignment sites (sCF) (Minh et al., 2020a). In addition to concordance factors, the values of the two discordance factors for each branch provide information about the relative contribution of stochastic error or more complex evolutionary processes to the signal, such as introgression. Highly uneven discordance factors are an indication of introgression, while discordance factors that are very similar to their concordance factors are indicative of substantial phylogenetic error or incomplete lineage sorting (Huson et al., 2005). We estimated gCFs using the gene trees and sCFs using the concatenated alignment, and repeated the analysis in the case that multiple phylogenetic resolutions were identified using IQ-TREE and ASTRAL-III.

The reconstructed species tree from ASTRAL-III was used as input for a Bayesian molecular dating analysis. We used highly efficient Bayesian dating using approximate likelihood computation (Thorne et al., 1998) as implemented in MCMCtree in PAML v4.8 (Yang, 2007). To reduce violation of the most common tree priors for molecular dating (Angelis & Dos Reis, 2015) and the impact of gene tree discordance on substitution rate estimates (Mendes & Hahn, 2016), we only included genomic regions with gene trees concordant with the species tree, in addition to other forms of filtering. Acceptable loci were partitioned by codon position, each modelled under a GTR+ Γ substitution model. We used an uncorrelated gamma prior on rates across lineages and a birth-death prior for divergence times. The posterior distribution was sampled using Markov chain Monte Carlo (MCMC) every $1x10^3$ steps over $1x10^7$ steps, after a burn-in phase of $1x10^6$ steps. We verified convergence to the stationary distribution by comparing the results from two independent runs, and confirming that the effective sample sizes for all parameters were above 1000 using the R package coda (Plummer et al., 2006).

Absolute times of divergence were estimated using a time-calibration strategy focusing on the deepest nodes of the tree (Duchêne et al., 2014). Four fossil calibrations were included, all based on nodes with consistently strong phylogenetic support in previous studies and using routinely-used fossils with robust taxonomic placement (e.g. Law et al., 2018). The split between Caniformes and Feliformes was calibrated using a conservative range for the age of the oldest known fossil of family Viverravidae (65 – 50 Mya; Benton et al., 2015; Fox et al., 2010; Wang & Tedford, 2008). The split between Canidae and other Caniformes was calibrated using a fossil of the genus *Amphicticeps* (32.8 – 30.4 Mya; Wang et al., 2005). The split between Pinnipedia and Musteloidea was calibrated using fossils of the stem musteloid of the genus *Mustelictis* (32.8 – 23.3 Mya; Rybczynski et al., 2009; Wang et al., 2005). To maximise the quality of our estimates of molecular rates and dates, we also calibrated the timing of the split between the pinnipeds *Odobenus rosmarus* and *Mirounga angustirostris*, using a fossil of the genus *Proneotherium* (20.4 – 13.8 Mya; Deméré et al., 2003). All fossil calibrations were set to follow a uniform distribution with soft maximum bounds.

#### *Nucleotide diversity*

For the sable, analysis of the heterozygosity distribution along chromosomes revealed that the ends of many pseudochromosomes had nearly double the heterozygosity than the median (> 2.5 SNPs per kbp vs median of 1.44 SNPs per kbp; Figure 2B, dark red color), while at least eight of the pseudochromosomes had very long stretches of low diversity (Figure 2B, blue color) that might be considered as runs of homozygosity (ROHs). Such distributed ROHs are often interpreted to be a consequence of recent inbreeding (Ceballos et al., 2018). This sable was sampled in the Greater Khingan mountains (Heilongjiang Province, China), very close to the edge of the species' range (Liu et al., 2020). The usual decline in abundance at the periphery, coupled with partial isolation from the main part of the species' range, might have resulted in recent inbreeding, which could explain the observed ROHs. Such a positive relationship between intraspecific SV counts and census (population) size has also been demonstrated for other species (Weissensteiner et al., 2020).

The pseudochromosome X (*ps_chrX)* of sable, had a uniform full (relative) coverage (0.75x - 1.25x relative to median whole genome coverage, SI Figure S5B), similar to the wolverine individual (SI Figure S5C). In contrast, we observed a clearly visible pseudoautosomal region (PAR) in the male tayra individual (SI Figure S5A), which had full (relative) coverage, whereas the rest of ps_chrX had half coverage. Additional verification using the SRY protein gene sequence as a Y-specific marker confirmed this result. The orthologous full-length CDS of SRY was detected only in the tayra assembly, while only partial hits with low similarity were observed in the two other assemblies. This provides evidence that the sable individual is female, not male, not XXY (Klinefelter syndrome in humans; Wikström & Dunkel, 2011), nor XX with translocation of the SRY locus to the X chromosome (de la Chapelle syndrome in humans; De la Chapelle et al., 1972).

#### *Structural variants*

Prior to SV calling, tayra reads were downsampled to ~38x using seqtk v1.3 (https://github.com/lh3/seqtk) to match the total coverage of wolverine and sable libraries so as to avoid bias in variant calling. Three SV callers, Manta v1.6.0 (Chen et al., 2016), Whamg v1.7.0 (Kronenberg et al., 2015) and Lumpy v0.2.13 (Layer et al., 2014) were selected based on their sensitivity and precision (Cameron et al., 2019), and used to identify putative SV events in the three Guloninae genomes. Manta combines paired-read (PR) and split-read (SR) evidence during SV discovery, along with SV breakend assembly (AS) to base-pair resolution. Whamg implements PR and SR support and Lumpy, a probabilistic CNV discovery tool, uses a combined approach of PR, SR and read-depth (RD) for SV detection. Intersecting results of multiple SV callers utilizing different methods for SV detection has previously been shown to improve accuracy of variant call sets (Kosugi et al., 2019; Pirooznia et al., 2015) although this approach is highly sensitive to the chosen set of callers (Cameron et al., 2019).

High coverage joint Manta variant calls (depth greater than 3x the median chromosome depth near one or both SV breakends) mainly caused by reads mapping in low complexity regions were filtered out as well as reads with MAPQ<30 for SV breakpoint support. We retained variant calls for which all samples passed all sample-level filters (FILTER=PASS), filtered out calls with genotype quality below 30 (GQ<30), and kept calls with paired-read (PR) and split-read (SR) support of PR>=3 and SR>=3. Unlocalized and unplaced scaffolds were removed and only scaffolds assigned to chromosomes were included in further analysis.

Whamg SV calls of size <50bp and >2Mb were filtered out to improve call accuracy, as well as calls with fewer than 10 supporting reads (total support, INFO field "A"). Calls with GQ<30 were filtered out. To reduce the number of false positive calls, we filtered out events flagged as "BND" with high cross-chromosomal mappings (CW>0.2), as Whamg is aware of but does not specifically call translocations. We further removed calls of all SV types with max(CW)<0.2 as these calls are associated with poorly mapped regions. We filtered out calls for which total evidence (paired and/or split reads) supporting the variant (FORMAT/SU field) was below 10 (SU<10).

Both Whamg and Lumpy SV call sets were genotyped with Svtyper v0.7.1 (Chiang et al., 2015) prior to filtering for genotype quality. Survivor v1.0.7 (Jeffares et al., 2017) was used to merge and compare SV call sets obtained from the three SV caller methods within and among samples. For each species and library insert size, we first merged SV events of the same type, called by at least two SV callers, with start/end positions detected within +/-1000 bp. We then intersected SV calls among the three species to obtain SVs private for each species (species-specific) and shared among all three species. To filter out SV calls overlapping gaps in the reference genome assemblies, we removed all SV calls overlapping stretches of "Ns" longer than 30 bases.

To annotate the final SV set, we first mapped annotations in gff3 format with Liftoff v1.5.1 (Shumate & Salzberg, 2020) between the domestic ferret draft genome assembly (MusPutFur1.0, GCF_000215625.1) and the chromosome-length domestic ferret assembly (DNA Zoo), which were used as a reference. We then reordered parent and children features with gff3sort (Zhu et al., 2017). Ensembl Variant Effect Predictor v101.0 (McLaren et al., 2016) was used to annotate variants affecting protein-coding genes with the maximum SV size set to 200 Mb. Annotated SV sets were further filtered for species-specific variants and SV type. Functional and biological roles of genes affected by SVs were explored using literature sources and online databases, including OrthoDB v10 (Kriventseva et al., 2019), Uniprot (The UniProt Consortium, 2017), and NCBI Entrez Gene (Maglott et al., 2011). Gene ontology analysis was performed with Shiny GO (Ge et al., 2020) with an FDR < 0.05 for each SV (excluding inversions) overlapping multiple protein-coding genes.

Supplemental Tables S4-S6 - References

Amos, J. S., Huang, L., Thevenon, J., Kariminedjad, A., Beaulieu, C. L., Masurel-Paulet, A., Najmabadi, H., Fattahi, Z., Beheshtian, M., Tonekaboni, S. H., Tang, S., Helbig, K. L., Alcaraz, W., Rivière, J.-B., Faivre, L., Innes, A. M., Lebel, R. R., Boycott, K. M., & Care4Rare Canada Consortium. (2017). Autosomal recessive mutations in THOC6 cause intellectual disability: syndrome delineation requiring forward and reverse phenotyping. *Clinical Genetics*, *91*(1), 92–99.

Avellino, R., Carrella, S., Pirozzi, M., Risolino, M., Salierno, F. G., Franco, P., Stoppelli, P., Verde, P., Banfi, S., & Conte, I. (2013). miR-204 targeting of Ankrd13A controls both mesenchymal neural crest and lens cell migration. *PloS One*, *8*(4), e61099.

Avnet, S., Lemma, S., Errani, C., Falzetti, L., Panza, E., Columbaro, M., Nanni, C., & Baldini, N. (2020). Benign albeit glycolytic: MCT4 expression and lactate release in giant cell tumour of bone. *Bone*, *134*, 115302.

Bähler, M., & Rhoads, A. (2002). Calmodulin signaling via the IQ motif. *FEBS Letters*, *513*(1), 107–113.

Banci, L., Bertini, I., Ciofi-Baffoni, S., Jaiswal, D., Neri, S., Peruzzini, R., & Winkelmann, J. (2012). Structural characterization of CHCHD5 and CHCHD7: two atypical human twin CX9C proteins. *Journal of Structural Biology*, *180*(1), 190–200.

Bao, J., Zhang, J., Zheng, H., Xu, C., & Yan, W. (2010). UBQLN1 interacts with SPEM1 and participates in spermiogenesis. *Molecular and Cellular Endocrinology*, *327*(1-2), 89–97.

Bauer, A., De Lucia, M., Jagannathan, V., Mezzalira, G., Casal, M. L., Welle, M. M., & Leeb, T. (2017). A Large Deletion in the NSDHL Gene in Labrador Retrievers with a Congenital Cornification Disorder. *G3* , *7*(9), 3115–3121.

Baumann, P., Schriever, S. C., Kullmann, S., Zimprich, A., Feuchtinger, A., Amarie, O., Peter, A., Walch, A., Gailus-Durner, V., Fuchs, H., Hrabě de Angelis, M., Wurst, W., Tschöp, M. H., Heni, M., Hölter, S. M., & Pfluger, P. T. (2019). Dusp8 affects hippocampal size and behavior in mice and humans. *Scientific Reports*, *9*(1), 19483.

Belizaire, R., Komanduri, C., Wooten, K., Chen, M., Thaller, C., & Janz, R. (2004). Characterization of synaptogyrin 3 as a new synaptic vesicle protein. *The Journal of Comparative Neurology*, *470*(3), 266–281.

Blume, M., Inoguchi, F., Sugiyama, T., Owada, Y., Osumi, N., Aimi, Y., Taki, K., & Katsuyama, Y. (2017). Dab1 contributes differently to the morphogenesis of the hippocampal subdivisions. *Development, Growth & Differentiation*, *59*(8), 657–673.

Browning, A. C., Figueiredo, G. S., Baylis, O., Montgomery, E., Beesley, C., Molinari, E., Figueiredo, F. C., & Sayer, J. A. (2019). A case of ocular cystinosis associated with two potentially severe CTNS mutations. *Ophthalmic Genetics*, *40*(2), 157–160.

Brown, J. A., Eberhardt, D. M., Schrick, F. N., Roberts, M. P., & Godkin, J. D. (2003). Expression of retinol-binding protein and cellular retinol-binding protein in the bovine ovary. *Molecular Reproduction and Development*, *64*(3), 261–269.

Cabral, W. A., Ishikawa, M., Garten, M., Makareeva, E. N., Sargent, B. M., Weis, M., Barnes, A. M.,

Webb, E. A., Shaw, N. J., Ala-Kokko, L., Lacbawan, F. L., Högler, W., Leikin, S., Blank, P. S., Zimmerberg, J., Eyre, D. R., Yamada, Y., & Marini, J. C. (2016). Absence of the ER Cation Channel TMEM38B/TRIC-B Disrupts Intracellular Calcium Homeostasis and Dysregulates Collagen Synthesis in Recessive Osteogenesis Imperfecta. *PLoS Genetics*, *12*(7), e1006156.

Chen, D., Teng, J. M., North, P. E., Lapinski, P. E., & King, P. D. (2019). RASA1-dependent cellular export of collagen IV controls blood and lymphatic vascular development. *The Journal of Clinical Investigation*, *129*(9), 3545–3561.

Chen, Q., Denard, B., Lee, C.-E., Han, S., Ye, J. S., & Ye, J. (2016). Inverting the Topology of a Transmembrane Protein by Regulating the Translocation of the First Transmembrane Helix. *Molecular Cell*, *63*(4), 567–578.

Cheong, A., Lingutla, R., & Mager, J. (2020). Expression analysis of mammalian mitochondrial ribosomal protein genes. *Gene Expression Patterns: GEP*, *38*, 119147.

Choi, Y. J., Kim, S., Choi, Y., Nielsen, T. B., Yan, J., Lu, A., Ruan, J., Lee, H.-R., Wu, H., Spellberg, B., & Jung, J. U. (2019). SERPINB1-mediated checkpoint of inflammatory caspase activation. *Nature Immunology*, *20*(3), 276–287.

Coan, P. M., Vaughan, O. R., Sekita, Y., Finn, S. L., Burton, G. J., Constancia, M., & Fowden, A. L. (2010). Adaptations in placental phenotype support fetal growth during undernutrition of pregnant mice. *The Journal of Physiology*, *588*(Pt 3), 527–538.

Crocco, P., Saiardi, A., Wilson, M. S., Maletta, R., Bruni, A. C., Passarino, G., & Rose, G. (2016). Contribution of polymorphic variation of inositol hexakisphosphate kinase 3 (IP6K3) gene promoter to the susceptibility to late onset Alzheimer's disease. *Biochimica et Biophysica Acta*, *1862*(9), 1766–1773.

D'Ambrosio, D. N., Clugston, R. D., & Blaner, W. S. (2011). Vitamin A metabolism: an update. *Nutrients*, *3*(1), 63–103.

Davydova, E., Ho, A. Y. Y., Malecki, J., Moen, A., Enserink, J. M., Jakobsson, M. E., Loenarz, C., & Falnes, P. Ø. (2014). Identification and characterization of a novel evolutionarily conserved lysine-specific methyltransferase targeting eukaryotic translation elongation factor 2 (eEF2). *The Journal of Biological Chemistry*, *289*(44), 30499–30510.

Del Dotto, V., Fogazza, M., Musiani, F., Maresca, A., Aleo, S. J., Caporali, L., La Morgia, C., Nolli, C., Lodi, T., Goffrini, P., Chan, D., Carelli, V., Rugolo, M., Baruffini, E., & Zanna, C. (2018). Deciphering OPA1 mutations pathogenicity by combined analysis of human, mouse and yeast cell models. *Biochimica et Biophysica Acta, Molecular Basis of Disease*, *1864*(10), 3496–3514.

Dong, F., Shinohara, K., Botilde, Y., Nabeshima, R., Asai, Y., Fukumoto, A., Hasegawa, T., Matsuo, M., Takeda, H., Shiratori, H., Nakamura, T., & Hamada, H. (2014). Pih1d3 is required for cytoplasmic preassembly of axonemal dynein in mouse sperm. *The Journal of Cell Biology*, *204*(2), 203–213.

Dreiza, C. M., Komalavilas, P., Furnish, E. J., Flynn, C. R., Sheller, M. R., Smoke, C. C., Lopes, L. B., & Brophy, C. M. (2010). The small heat shock protein, HSPB6, in muscle function and disease. *Cell Stress & Chaperones*, *15*(1), 1–11.

Feng, W., & Zhang, M. (2009). Organization and dynamics of PDZ-domain-related supramodules in the postsynaptic density. *Nature Reviews. Neuroscience*, *10*(2), 87–99.

Feng, Y., Xu, X., Zhang, Y., Ding, J., Wang, Y., Zhang, X., Wu, Z., Kang, L., Liang, Y., Zhou, L., Song, S., Zhao, K., & Ye, Q. (2015). HPIP is upregulated in colorectal cancer and regulates colorectal cancer cell proliferation, apoptosis and invasion. *Scientific Reports*, *5*, 9429.

Geisinger, A., Alsheimer, M., Baier, A., Benavente, R., & Wettstein, R. (2005). The mammalian gene pecanex 1 is differentially expressed during spermatogenesis. *Biochimica et Biophysica Acta*, *1728*(1-2), 34–43.

Gerits, N., Van Belle, W., & Moens, U. (2007). Transgenic mice expressing constitutive active MAPKAPK5 display gender-dependent differences in exploration and activity. *Behavioral and Brain Functions: BBF*, *3*, 58.

Gòdia, M., Casellas, J., Ruiz-Herrera, A., Rodríguez-Gil, J. E., Castelló, A., Sánchez, A., & Clop, A. (2020). Whole genome sequencing identifies allelic ratio distortion in sperm involving genes related to spermatogenesis in a swine model. *DNA Research: An International Journal for Rapid*

*Publication of Reports on Genes and Genomes*, *27*(5).
Goldfarb, K. C., & Cech, T. R. (2017). Targeted CRISPR disruption reveals a role for RNase MRP RNA in human preribosomal RNA processing. *Genes & Development*, *31*(1), 59–71.
Grayson, P., & Civetta, A. (2012). Positive Selection and the Evolution of izumo Genes in Mammals. *International Journal of Evolutionary Biology*, *2012*, 958164.
Griffiths, M. R., Botto, M., Morgan, B. P., Neal, J. W., & Gasque, P. (2018). CD93 regulates central nervous system inflammation in two mouse models of autoimmune encephalomyelitis. *Immunology*, *155*(3), 346–355.
Guo, H. X., Cun, W., Liu, L. D., Dong, S. Z., Wang, L. C., Dong, C. H., & Li, Q. H. (2006). Protein encoded by HSV-1 stimulation-related gene 1 (HSRG1) interacts with and inhibits SV40 large T antigen. *Cell Proliferation*, *39*(6), 507–518.
He, Y.-W., Li, H., Zhang, J., Hsu, C.-L., Lin, E., Zhang, N., Guo, J., Forbush, K. A., & Bevan, M. J. (2004). The extracellular matrix protein mindin is a pattern-recognition molecule for microbial pathogens. *Nature Immunology*, *5*(1), 88–97.
Hill, M., Pařízek, A., Cibula, D., Kancheva, R., Jirásek, J. E., Jirkovská, M., Velíková, M., Kubátová, J., Klímková, M., Pašková, A., Zižka, Z., Kancheva, L., Kazihnitková, H., Zamrazilová, L., & Stárka, L. (2010). Steroid metabolome in fetal and maternal body fluids in human late pregnancy. *The Journal of Steroid Biochemistry and Molecular Biology*, *122*(4), 114–132.
Hnia, K., Tronchère, H., Tomczak, K. K., Amoasii, L., Schultz, P., Beggs, A. H., Payrastre, B., Mandel, J. L., & Laporte, J. (2011). Myotubularin controls desmin intermediate filament architecture and mitochondrial dynamics in human and mouse skeletal muscle. *The Journal of Clinical Investigation*, *121*(1), 70–85.
Horibata, Y., Elpeleg, O., Eran, A., Hirabayashi, Y., Savitzki, D., Tal, G., Mandel, H., & Sugimoto, H. (2018). EPT1 (selenoprotein I) is critical for the neural development and maintenance of plasmalogen in humans. *Journal of Lipid Research*, *59*(6), 1015–1026.
Im, S.-H., Kim, S.-H., Azam, T., Venkatesh, N., Dinarello, C. A., Fuchs, S., & Souroujon, M. C. (2002). Rat interleukin-18 binding protein: cloning, expression, and characterization. *Journal of Interferon & Cytokine Research: The Official Journal of the International Society for Interferon and Cytokine Research*, *22*(3), 321–328.
Inoue, K., Maeda, N., Mori, T., Sekimoto, R., Tsushima, Y., Matsuda, K., Yamaoka, M., Suganami, T., Nishizawa, H., Ogawa, Y., Funahashi, T., & Shimomura, I. (2014). Possible involvement of Opa-interacting protein 5 in adipose proliferation and obesity. *PloS One*, *9*(2), e87661.
Ishizawa, T., Nozaki, Y., Ueda, T., & Takeuchi, N. (2008). The human mitochondrial translation release factor HMRF1L is methylated in the GGQ motif by the methyltransferase HMPrmC. *Biochemical and Biophysical Research Communications*, *373*(1), 99–103.
Jakobsson, T., Venteclef, N., Toresson, G., Damdimopoulos, A. E., Ehrlund, A., Lou, X., Sanyal, S., Steffensen, K. R., Gustafsson, J.-A., & Treuter, E. (2009). GPS2 is required for cholesterol efflux by triggering histone demethylation, LXR recruitment, and coregulator assembly at the ABCG1 locus. *Molecular Cell*, *34*(4), 510–518.
Johnson, R. J., Stenvinkel, P., Andrews, P., Sánchez-Lozada, L. G., Nakagawa, T., Gaucher, E., Andres-Hernando, A., Rodriguez-Iturbe, B., Jimenez, C. R., Garcia, G., Kang, D.-H., Tolan, D. R., & Lanaspa, M. A. (2020). Fructose metabolism as a common evolutionary pathway of survival associated with climate change, food shortage and droughts. *Journal of Internal Medicine*, *287*(3), 252–262.
Kang, C. B., Hong, Y., Dhe-Paganon, S., & Yoon, H. S. (2008). FKBP family proteins: immunophilins with versatile biological functions. *Neuro-Signals*, *16*(4), 318–325.
Katzen, J., Wagner, B. D., Venosa, A., Kopp, M., Tomer, Y., Russo, S. J., Headen, A. C., Basil, M. C., Stark, J. M., Mulugeta, S., Deterding, R. R., & Beers, M. F. (2019). An SFTPC BRICHOS mutant links epithelial ER stress and spontaneous lung fibrosis. *JCI Insight*, *4*(6). https://doi.org/10.1172/jci.insight.126125
Kazmierczak, M., Harris, S. L., Kazmierczak, P., Shah, P., Starovoytov, V., Ohlemiller, K. K., & Schwander, M. (2015). Progressive Hearing Loss in Mice Carrying a Mutation in Usp53. *The Journal of Neuroscience: The Official Journal of the Society for Neuroscience*, *35*(47),

15582–15598.
Kennell, J. A., Richards, N. W., Schaner, P. E., & Gumucio, D. L. (2001). cDNA cloning, chromosomal localization and evolutionary analysis of mouse vacuolar ATPase subunit D, Atp6m. *Cytogenetics and Cell Genetics*, *92*(3-4), 337–341.
Kielkowski, P., Buchsbaum, I. Y., Kirsch, V. C., Bach, N. C., Drukker, M., Cappello, S., & Sieber, S. A. (2020). FICD activity and AMPylation remodelling modulate human neurogenesis. *Nature Communications*, *11*(1), 517.
Kim, J. H., Gurumurthy, C. B., Band, H., & Band, V. (2010). Biochemical characterization of human Ecdysoneless reveals a role in transcriptional regulation. *Biological Chemistry*, *391*(1), 9–19.
Kim, J., Ishiguro, K.-I., Nambu, A., Akiyoshi, B., Yokobayashi, S., Kagami, A., Ishiguro, T., Pendas, A. M., Takeda, N., Sakakibara, Y., Kitajima, T. S., Tanno, Y., Sakuno, T., & Watanabe, Y. (2015). Meikin is a conserved regulator of meiosis-I-specific kinetochore function. *Nature*, *517*(7535), 466–471.
Kim, M. Y., Lee, H. K., Park, J. S., Park, S. H., Kwon, H. B., & Soh, J. (1999). Identification of a zeta-crystallin (quinone reductase)-like 1 gene (CRYZL1) mapped to human chromosome 21q22.1. *Genomics*, *57*(1), 156–159.
Kitchener, A. C., Meloro, C., & Williams, T. M. (2018). Form and function of the musteloids. In *Biology and Conservation of Musteloids*. Oxford University Press.
Koc, E. C., Burkhart, W., Blackburn, K., Moseley, A., Koc, H., & Spremulli, L. L. (2000). A Proteomics Approach to the Identification of Mammalian Mitochondrial Small Subunit Ribosomal Proteins *. *The Journal of Biological Chemistry*, *275*(42), 32585–32591.
Kohroki, J., Nishiyama, T., Nakamura, T., & Masuho, Y. (2005). ASB proteins interact with Cullin5 and Rbx2 to form E3 ubiquitin ligase complexes. *FEBS Letters*, *579*(30), 6796–6802.
Kriventseva, E. V., Kuznetsov, D., Tegenfeldt, F., Manni, M., Dias, R., Simão, F. A., & Zdobnov, E. M. (2019). OrthoDB v10: sampling the diversity of animal, plant, fungal, protist, bacterial and viral genomes for evolutionary and functional annotations of orthologs. *Nucleic Acids Research*, *47*(D1), D807–D811.
Kuo, P.-L., Chiang, H.-S., Wang, Y.-Y., Kuo, Y.-C., Chen, M.-F., Yu, I.-S., Teng, Y.-N., Lin, S.-W., & Lin, Y.-H. (2013). SEPT12-microtubule complexes are required for sperm head and tail formation. *International Journal of Molecular Sciences*, *14*(11), 22102–22116.
Kuriakose, T., & Kanneganti, T.-D. (2018). ZBP1: Innate Sensor Regulating Cell Death and Inflammation. *Trends in Immunology*, *39*(2), 123–134.
Lal-Nag, M., & Morin, P. J. (2009). The claudins. *Genome Biology*, *10*(8), 235.
Larsen, K., Momeni, J., Farajzadeh, L., & Callesen, H. (2017). Splice variants of porcine PPHLN1 encoding periphilin-1. *Gene Reports*, *7*, 176–183.
Lei, Y., Wen, H., Yu, Y., Taxman, D. J., Zhang, L., Widman, D. G., Swanson, K. V., Wen, K.-W., Damania, B., Moore, C. B., Giguère, P. M., Siderovski, D. P., Hiscott, J., Razani, B., Semenkovich, C. F., Chen, X., & Ting, J. P.-Y. (2012). The mitochondrial proteins NLRX1 and TUFM form a complex that regulates type I interferon and autophagy. *Immunity*, *36*(6), 933–946.
Li, Q., Korzan, W. J., Ferrero, D. M., Chang, R. B., Roy, D. S., Buchi, M., Lemon, J. K., Kaur, A. W., Stowers, L., Fendt, M., & Liberles, S. D. (2013). Synchronous evolution of an odor biosynthesis pathway and behavioral response. *Current Biology: CB*, *23*(1), 11–20.
Liu, S., Gong, X., Yan, X., Peng, T., Baker, J. C., Li, L., Robben, P. M., Ravindran, S., Andersson, L. A., Cole, A. B., & Roche, T. E. (2001). Reaction mechanism for mammalian pyruvate dehydrogenase using natural lipoyl domain substrates. *Archives of Biochemistry and Biophysics*, *386*(2), 123–135.
Luczkowska, K., Stekelenburg, C., Sloan-Béna, F., Ranza, E., Gastaldi, G., Schwitzgebel, V., & Maechler, P. (2020). Hyperinsulinism associated with GLUD1 mutation: allosteric regulation and functional characterization of p.G446V glutamate dehydrogenase. *Human Genomics*, *14*(1), 9.
Lupo, G., Nisi, P. S., Esteve, P., Paul, Y.-L., Novo, C. L., Sidders, B., Khan, M. A., Biagioni, S., Liu, H.-K., Bovolenta, P., Cacci, E., & Rugg-Gunn, P. J. (2018). Molecular profiling of aged neural progenitors identifies Dbx2 as a candidate regulator of age-associated neurogenic decline. *Aging Cell*, *17*(3), e12745.

Maglott, D., Ostell, J., Pruitt, K. D., & Tatusova, T. (2011). Entrez Gene: gene-centered information at NCBI. *Nucleic Acids Research*, *39*(Database issue), D52–D57.

Makarova, O. V., Makarov, E. M., & Lührmann, R. (2001). The 65 and 110 kDa SR-related proteins of the U4/U6.U5 tri-snRNP are essential for the assembly of mature spliceosomes. *The EMBO Journal*, *20*(10), 2553–2563.

Martinez-Duncker, I., Dupré, T., Piller, V., Piller, F., Candelier, J.-J., Trichet, C., Tchernia, G., Oriol, R., & Mollicone, R. (2005). Genetic complementation reveals a novel human congenital disorder of glycosylation of type II, due to inactivation of the Golgi CMP-sialic acid transporter. *Blood*, *105*(7), 2671–2676.

Masure, S., Cik, M., Hoefnagel, E., Nosrat, C. A., Van der Linden, I., Scott, R., Van Gompel, P., Lesage, A. S., Verhasselt, P., Ibáñez, C. F., & Gordon, R. D. (2000). Mammalian GFRalpha -4, a divergent member of the GFRalpha family of coreceptors for glial cell line-derived neurotrophic factor family ligands, is a receptor for the neurotrophic factor persephin. *The Journal of Biological Chemistry*, *275*(50), 39427–39434.

Matsubara, T., Kokabu, S., Nakatomi, C., Kinbara, M., Maeda, T., Yoshizawa, M., Yasuda, H., Takano-Yamamoto, T., Baron, R., & Jimi, E. (2018). The Actin-Binding Protein PPP1r18 Regulates Maturation, Actin Organization, and Bone Resorption Activity of Osteoclasts. *Molecular and Cellular Biology*, *38*(4).

McDaniel, P., & Wu, X. (2009). Identification of oocyte-selective NLRP genes in rhesus macaque monkeys (Macaca mulatta). *Molecular Reproduction and Development*, *76*(2), 151–159.

Metzger, J., Karwath, M., Tonda, R., Beltran, S., Águeda, L., Gut, M., Gut, I. G., & Distl, O. (2015). Runs of homozygosity reveal signatures of positive selection for reproduction traits in breed and non-breed horses. *BMC Genomics*, *16*, 764.

Mironova, E., & Millette, C. F. (2008). Expression of the diaphanous-related formin proteins mDia1 and mDia2 in the rat testis. *Developmental Dynamics: An Official Publication of the American Association of Anatomists*, *237*(8), 2170–2176.

Mizuarai, S., Miki, S., Araki, H., Takahashi, K., & Kotani, H. (2005). Identification of dicarboxylate carrier Slc25a10 as malate transporter in de novo fatty acid synthesis. *The Journal of Biological Chemistry*, *280*(37), 32434–32441.

Montanez, E., Wickström, S. A., Altstätter, J., Chu, H., & Fässler, R. (2009). Alpha-parvin controls vascular mural cell recruitment to vessel wall by regulating RhoA/ROCK signalling. *The EMBO Journal*, *28*(20), 3132–3144.

Nakamura, S., Kahyo, T., Tao, H., Shibata, K., Kurabe, N., Yamada, H., Shinmura, K., Ohnishi, K., & Sugimura, H. (2015). Novel roles for LIX1L in promoting cancer cell proliferation through ROS1-mediated LIX1L phosphorylation. *Scientific Reports*, *5*, 13474.

Nelson, L., Anderson, S., Archibald, A. L., Rhind, S., Lu, Z. H., Condie, A., McIntyre, N., Thompson, J., Nenutil, R., Vojtesek, B., Whitelaw, C. B. A., Little, T. J., & Hupp, T. (2008). An animal model to evaluate the function and regulation of the adaptively evolving stress protein SEP53 in oesophageal bile damage responses. *Cell Stress & Chaperones*, *13*(3), 375–385.

Nomura, Y., Roston, D., Montemayor, E. J., Cui, Q., & Butcher, S. E. (2018). Structural and mechanistic basis for preferential deadenylation of U6 snRNA by Usb1. *Nucleic Acids Research*, *46*(21), 11488–11501.

Papadopoulos, C., Kirchner, P., Bug, M., Grum, D., Koerver, L., Schulze, N., Poehler, R., Dressler, A., Fengler, S., Arhzaouy, K., Lux, V., Ehrmann, M., Weihl, C. C., & Meyer, H. (2017). VCP/p97 cooperates with YOD1, UBXD1 and PLAA to drive clearance of ruptured lysosomes by autophagy. *The EMBO Journal*, *36*(2), 135–150.

Park, K. M., Kang, E., Jeon, Y.-J., Kim, N., Kim, N.-S., Yoo, H.-S., Yeom, Y. I., & Kim, S. J. (2007). Identification of novel regulators of apoptosis using a high-throughput cell-based screen. *Molecules and Cells*, *23*(2), 170–174.

Paschen, S. A., Rothbauer, U., Káldi, K., Bauer, M. F., Neupert, W., & Brunner, M. (2000). The role of the TIM8-13 complex in the import of Tim23 into mitochondria. *The EMBO Journal*, *19*(23), 6392–6400.

Pei, J., & Grishin, N. V. (2012). Unexpected diversity in Shisa-like proteins suggests the importance of

their roles as transmembrane adaptors. *Cellular Signalling*, *24*(3), 758–769.
Perumal, K., Sinha, K., Henning, D., & Reddy, R. (2001). Purification, characterization, and cloning of the cDNA of human signal recognition particle RNA 3'-adenylating enzyme. *The Journal of Biological Chemistry*, *276*(24), 21791–21796.
Picollo, A., & Pusch, M. (2005). Chloride/proton antiporter activity of mammalian CLC proteins ClC-4 and ClC-5. *Nature*, *436*(7049), 420–423.
Pierre, K., Parent, A., Jayet, P.-Y., Halestrap, A. P., Scherrer, U., & Pellerin, L. (2007). Enhanced expression of three monocarboxylate transporter isoforms in the brain of obese mice. *The Journal of Physiology*, *583*(Pt 2), 469–486.
Premzl, M. (2016). Comparative genomic analysis of eutherian tumor necrosis factor ligand genes. *Immunogenetics*, *68*(2), 125–132.
Pu, J., Schindler, C., Jia, R., Jarnik, M., Backlund, P., & Bonifacino, J. S. (2015). BORC, a multisubunit complex that regulates lysosome positioning. *Developmental Cell*, *33*(2), 176–188.
Qureshi, T., Bjørkmo, M., Nordengen, K., Gundersen, V., Utheim, T. P., Watne, L. O., Storm-Mathisen, J., Hassel, B., & Chaudhry, F. A. (2020). Slc38a1 Conveys Astroglia-Derived Glutamine into GABAergic Interneurons for Neurotransmitter GABA Synthesis. *Cells* , *9*(7). https://doi.org/10.3390/cells9071686
Raffaello, A., De Stefani, D., Sabbadin, D., Teardo, E., Merli, G., Picard, A., Checchetto, V., Moro, S., Szabò, I., & Rizzuto, R. (2013). The mitochondrial calcium uniporter is a multimer that can include a dominant-negative pore-forming subunit. *The EMBO Journal*, *32*(17), 2362–2376.
Rauniyar, K., Jha, S. K., & Jeltsch, M. (2018). Biology of Vascular Endothelial Growth Factor C in the Morphogenesis of Lymphatic Vessels. *Frontiers in Bioengineering and Biotechnology*, *6*, 7.
Reynolds, A., Qiao, H., Yang, Y., Chen, J. K., Jackson, N., Biswas, K., Holloway, J. K., Baudat, F., de Massy, B., Wang, J., Höög, C., Cohen, P. E., & Hunter, N. (2013). RNF212 is a dosage-sensitive regulator of crossing-over during mammalian meiosis. *Nature Genetics*, *45*(3), 269–278.
Rismanchi, N., Soderblom, C., Stadler, J., Zhu, P.-P., & Blackstone, C. (2008). Atlastin GTPases are required for Golgi apparatus and ER morphogenesis. *Human Molecular Genetics*, *17*(11), 1591–1604.
Rivas, M. A., Graham, D., Sulem, P., Stevens, C., Desch, A. N., Goyette, P., Gudbjartsson, D., Jonsdottir, I., Thorsteinsdottir, U., Degenhardt, F., Mucha, S., Kurki, M. I., Li, D., D'Amato, M., Annese, V., Vermeire, S., Weersma, R. K., Halfvarson, J., Paavola-Sakki, P., … Wang, M. H. (2016). A protein-truncating R179X variant in RNF186 confers protection against ulcerative colitis. *Nature Communications*, *7*, 12342.
Salleron, L., Magistrelli, G., Mary, C., Fischer, N., Bairoch, A., & Lane, L. (2014). DERA is the human deoxyribose phosphate aldolase and is involved in stress response. *Biochimica et Biophysica Acta*, *1843*(12), 2913–2925.
Schlager, M. A., Kapitein, L. C., Grigoriev, I., Burzynski, G. M., Wulf, P. S., Keijzer, N., de Graaff, E., Fukuda, M., Shepherd, I. T., Akhmanova, A., & Hoogenraad, C. C. (2010). Pericentrosomal targeting of Rab6 secretory vesicles by Bicaudal-D-related protein 1 (BICDR-1) regulates neuritogenesis. *The EMBO Journal*, *29*(10), 1637–1651.
Shaheen, R., Jiang, N., Alzahrani, F., Ewida, N., Al-Sheddi, T., Alobeid, E., Musaev, D., Stanley, V., Hashem, M., Ibrahim, N., Abdulwahab, F., Alshenqiti, A., Sonmez, F. M., Saqati, N., Alzaidan, H., Al-Qattan, M. M., Al-Mohanna, F., Gleeson, J. G., & Alkuraya, F. S. (2019). Bi-allelic Mutations in FAM149B1 Cause Abnormal Primary Cilium and a Range of Ciliopathy Phenotypes in Humans. *American Journal of Human Genetics*, *104*(4), 731–737.
Shang, J., Xia, T., Han, Q.-Q., Zhao, X., Hu, M.-M., Shu, H.-B., & Guo, L. (2018). Quantitative Proteomics Identified TTC4 as a TBK1 Interactor and a Positive Regulator of SeV-Induced Innate Immunity. *Proteomics*, *18*(2).
Shiio, Y., Rose, D. W., Aur, R., Donohoe, S., Aebersold, R., & Eisenman, R. N. (2006). Identification and characterization of SAP25, a novel component of the mSin3 corepressor complex. *Molecular and Cellular Biology*, *26*(4), 1386–1397.
Shi, Y.-Q., Li, Y.-C., Hu, X.-Q., Liu, T., Liao, S.-Y., Guo, J., Huang, L., Hu, Z.-Y., Tang, A. Y. B., Lee, K.-F., Yeung, W. S. B., Han, C.-S., & Liu, Y.-X. (2009). Male germ cell-specific protein Trs4

binds to multiple proteins. *Biochemical and Biophysical Research Communications*, *388*(3), 583–588.

Shyu, M.-K., Lin, M.-C., Shih, J.-C., Lee, C.-N., Huang, J., Liao, C.-H., Huang, I.-F., Chen, H.-Y., Huang, M.-C., & Hsieh, F.-J. (2007). Mucin 15 is expressed in human placenta and suppresses invasion of trophoblast-like cells in vitro. *Human Reproduction* , *22*(10), 2723–2732.

Siepka, S. M., Yoo, S.-H., Park, J., Song, W., Kumar, V., Hu, Y., Lee, C., & Takahashi, J. S. (2007). Circadian mutant Overtime reveals F-box protein FBXL3 regulation of cryptochrome and period gene expression. *Cell*, *129*(5), 1011–1023.

Sillibourne, J. E., Delaval, B., Redick, S., Sinha, M., & Doxsey, S. J. (2007). Chromatin remodeling proteins interact with pericentrin to regulate centrosome integrity. *Molecular Biology of the Cell*, *18*(9), 3667–3680.

Singh, B. N., Gong, W., Das, S., Theisen, J. W. M., Sierra-Pagan, J. E., Yannopoulos, D., Skie, E., Shah, P., Garry, M. G., & Garry, D. J. (2019). Etv2 transcriptionally regulates Yes1 and promotes cell proliferation during embryogenesis. *Scientific Reports*, *9*(1), 9736.

Singh, P., Patel, R. K., Palmer, N., Grenier, J. K., Paduch, D., Kaldis, P., Grimson, A., & Schimenti, J. C. (2019). CDK2 kinase activity is a regulator of male germ cell fate. *Development*, *146*(21).

Sonna, L. A., Fujita, J., Gaffin, S. L., & Lilly, C. M. (2002). Invited review: Effects of heat and cold stress on mammalian gene expression. *Journal of Applied Physiology*, *92*(4), 1725–1742.

Sun-Wada, G. H., Murakami, H., Nakai, H., Wada, Y., & Futai, M. (2001). Mouse Atp6f, the gene encoding the 23-kDa proteolipid of vacuolar proton translocating ATPase. *Gene*, *274*(1-2), 93–99.

Tashita, C., Hoshi, M., Hirata, A., Nakamoto, K., Ando, T., Hattori, T., Yamamoto, Y., Tezuka, H., Tomita, H., Hara, A., & Saito, K. (2020). Kynurenine plays an immunosuppressive role in 2,4,6-trinitrobenzene sulfate-induced colitis in mice. *World Journal of Gastroenterology: WJG*, *26*(9), 918–932.

Tejada-Jiménez, M., Galván, A., & Fernández, E. (2011). Algae and humans share a molybdate transporter. *Proceedings of the National Academy of Sciences of the United States of America*, *108*(16), 6420–6425.

Tezuka, Y., Okada, M., Tada, Y., Yamauchi, J., Nishigori, H., & Sanbe, A. (2013). Regulation of neurite growth by inorganic pyrophosphatase 1 via JNK dephosphorylation. *PloS One*, *8*(4), e61649.

The UniProt Consortium. (2017). UniProt: the universal protein knowledgebase. *Nucleic Acids Research*, *45*(D1), D158–D169.

Touyama, K., Khan, M., Aoki, K., Matsuda, M., Hiura, F., Takakura, N., Matsubara, T., Harada, Y., Hirohashi, Y., Tamura, Y., Gao, J., Mori, K., Kokabu, S., Yasuda, H., Fujita, Y., Watanabe, K., Takahashi, Y., Maki, K., & Jimi, E. (2019). Bif-1/Endophilin B1/SH3GLB1 regulates bone homeostasis. *Journal of Cellular Biochemistry*, *120*(11), 18793–18804.

Tsantoulas, C., Denk, F., Signore, M., Nassar, M. A., Futai, K., & McMahon, S. B. (2018). Mice lacking Kcns1 in peripheral neurons show increased basal and neuropathic pain sensitivity. *Pain*, *159*(8), 1641–1651.

Valente, P., Romei, A., Fadda, M., Sterlini, B., Lonardoni, D., Forte, N., Fruscione, F., Castroflorio, E., Michetti, C., Giansante, G., Valtorta, F., Tsai, J.-W., Zara, F., Nieus, T., Corradi, A., Fassio, A., Baldelli, P., & Benfenati, F. (2019). Constitutive Inactivation of the PRRT2 Gene Alters Short-Term Synaptic Plasticity and Promotes Network Hyperexcitability in Hippocampal Neurons. *Cerebral Cortex* , *29*(5), 2010–2033.

Wehner, K. A., & Baserga, S. J. (2002). The sigma(70)-like motif: a eukaryotic RNA binding domain unique to a superfamily of proteins required for ribosome biogenesis. *Molecular Cell*, *9*(2), 329–339.

Weinstat-Saslow, D. L., Germino, G. G., Somlo, S., & Reeders, S. T. (1993). A transducin-like gene maps to the autosomal dominant polycystic kidney disease gene region. *Genomics*, *18*(3), 709–711.

Wild, T., Budzowska, M., Hellmuth, S., Eibes, S., Karemore, G., Barisic, M., Stemmann, O., & Choudhary, C. (2018). Deletion of APC7 or APC16 Allows Proliferation of Human Cells without the Spindle Assembly Checkpoint. *Cell Reports*, *25*(9), 2317–2328.e5.

Wiley, S. R., Cassiano, L., Lofton, T., Davis-Smith, T., Winkles, J. A., Lindner, V., Liu, H., Daniel, T. O., Smith, C. A., & Fanslow, W. C. (2001). A novel TNF receptor family member binds TWEAK and is implicated in angiogenesis. *Immunity*, *15*(5), 837–846.

Wu, S. Y., Thomas, M. C., Hou, S. Y., Likhite, V., & Chiang, C. M. (1999). Isolation of mouse TFIID and functional characterization of TBP and TFIID in mediating estrogen receptor and chromatin transcription. *The Journal of Biological Chemistry*, *274*(33), 23480–23490.

Wu, X., Quondamatteo, F., Lefever, T., Czuchra, A., Meyer, H., Chrostek, A., Paus, R., Langbein, L., & Brakebusch, C. (2006). Cdc42 controls progenitor cell differentiation and beta-catenin turnover in skin. *Genes & Development*, *20*(5), 571–585.

Xiao, Q., Wu, X.-L., Michal, J. J., Reeves, J. J., Busboom, J. R., Thorgaard, G. H., & Jiang, Z. (2006). A novel nuclear-encoded mitochondrial poly(A) polymerase PAPD1 is a potential candidate gene for the extreme obesity related phenotypes in mammals. *International Journal of Biological Sciences*, *2*(4), 171–178.

Xie, L., Qin, W.-X., He, X.-H., Shu, H.-Q., Yao, G.-F., Wan, D.-F., & Gu, J.-R. (2004). Differential gene expression in human hepatocellular carcinoma Hep3B cells induced by apoptosis-related gene BNIPL-2. *World Journal of Gastroenterology: WJG*, *10*(9), 1286–1291.

Xie, X.-K., Xu, Z.-K., Xu, K., & Xiao, Y.-X. (2020). DUSP19 mediates spinal cord injury-induced apoptosis and inflammation in mouse primary microglia cells via the NF-kB signaling pathway. *Neurological Research*, *42*(1), 31–38.

Xu, Z., Yang, L., Xu, S., Zhang, Z., & Cao, Y. (2015). The receptor proteins: pivotal roles in selective autophagy. *Acta Biochimica et Biophysica Sinica*, *47*(8), 571–580.

Yamakoshi, T., Makino, T., Ur Rehman, M., Yoshihisa, Y., Sugimori, M., & Shimizu, T. (2013). Trichohyalin-like 1 protein, a member of fused S100 proteins, is expressed in normal and pathologic human skin. *Biochemical and Biophysical Research Communications*, *432*(1), 66–72.

Yang, X., Matsuda, K., Bialek, P., Jacquot, S., Masuoka, H. C., Schinke, T., Li, L., Brancorsini, S., Sassone-Corsi, P., Townes, T. M., Hanauer, A., & Karsenty, G. (2004). ATF4 is a substrate of RSK2 and an essential regulator of osteoblast biology; implication for Coffin-Lowry Syndrome. *Cell*, *117*(3), 387–398.

Ye, W., Zhou, Y., Xu, B., Zhu, D., Rui, X., Xu, M., Shi, L., Zhang, D., & Jiang, J. (2019). CD247 expression is associated with differentiation and classification in ovarian cancer. *Medicine*, *98*(51), e18407.

Ye, Z., & Ting, J. P.-Y. (2008). NLR, the nucleotide-binding domain leucine-rich repeat containing gene family. *Current Opinion in Immunology*, *20*(1), 3–9.

Yockey, L. J., & Iwasaki, A. (2018). Interferons and Proinflammatory Cytokines in Pregnancy and Fetal Development. *Immunity*, *49*(3), 397–412.

Yu, Y.-H., Chang, Y.-C., Su, T.-H., Nong, J.-Y., Li, C.-C., & Chuang, L.-M. (2013). Prostaglandin reductase-3 negatively modulates adipogenesis through regulation of PPARγ activity. *Journal of Lipid Research*, *54*(9), 2391–2399.

Zhang, C., & Liang, Y. (2018). Latexin and hematopoiesis. *Current Opinion in Hematology*, *25*(4), 266–272.

Zhang, H., Ge, Y., He, P., Chen, X., Carina, A., Qiu, Y., Aga, D. S., & Ren, X. (2015). Interactive Effects of N6AMT1 and As3MT in Arsenic Biomethylation. *Toxicological Sciences: An Official Journal of the Society of Toxicology*, *146*(2), 354–362.

Zhang, W. J., & Wu, J. Y. (1998). Sip1, a novel RS domain-containing protein essential for pre-mRNA splicing. *Molecular and Cellular Biology*, *18*(2), 676–684.

Zhang, X., Azhar, G., Zhong, Y., & Wei, J. Y. (2006). Zipzap/p200 is a novel zinc finger protein contributing to cardiac gene regulation. *Biochemical and Biophysical Research Communications*, *346*(3), 794–801.

Zhao, H., Zhu, L., Zhu, Y., Cao, J., Li, S., Huang, Q., Xu, T., Huang, X., Yan, X., & Zhu, X. (2013). The Cep63 paralogue Deup1 enables massive de novo centriole biogenesis for vertebrate multiciliogenesis. *Nature Cell Biology*, *15*(12), 1434–1444.
